# Antigen-processing rewiring expose cryptic self promoting organ-specific autoimmunity

**DOI:** 10.64898/2025.12.12.693959

**Authors:** Astrid Brix Saksager, Sarah Rosenberg Asmussen, Freja Dahl Hede, Carolina Barra

**Affiliations:** Section for Bioinformatics, Department of Health Technology, Department of Health and Technology, Technical University of Denmark, 2800 Kongens Lyngby, Denmark

**Author notes:** **Corresponding Author:** Carolina Barra.

**Keywords:** autoimmunity, immunopeptidomics, antigen processing, mhc class II, autoantigens

## Abstract

Autoimmune disease is traditionally attributed to failure of immune tolerance toward normally presented self-antigens. Although strong HLA associations implicate antigen presentation in disease risk, whether altered antigen processing contributes directly to autoimmunity remains unclear. To address this question, we analysed experimentally verified human T-cell autoantigens across twenty autoimmune diseases and classified them according to their natural presentation on HLA-DR in healthy individuals as either tolerant (detectably presented on MHC) or cryptic (not detectably presented). These classes exhibited distinct molecular and clinical associations: cryptic autoantigens were enriched in membrane proteins with tissue-specific functions and were predominantly linked to organ-specific autoimmune diseases, whereas tolerant autoantigens were largely extracellular, enriched in immune-related functions, and associated with systemic autoimmunity. Analysis of immunopeptidomics datasets from rheumatoid arthritis and multiple sclerosis further revealed disease-associated shifts in peptide flanking residues and increased relative solvent accessibility compared with healthy donors, consistent with altered proteolytic processing. Cryptic proteins were markedly overrepresented among multiple sclerosis-derived ligands. Together, these findings define two mechanistically distinct routes to autoimmune activation: one driven by altered antigen processing that exposes previously unseen self-proteins, and another driven by breakdown of tolerance toward normally presented self-antigens.

**Graphical abstract:** 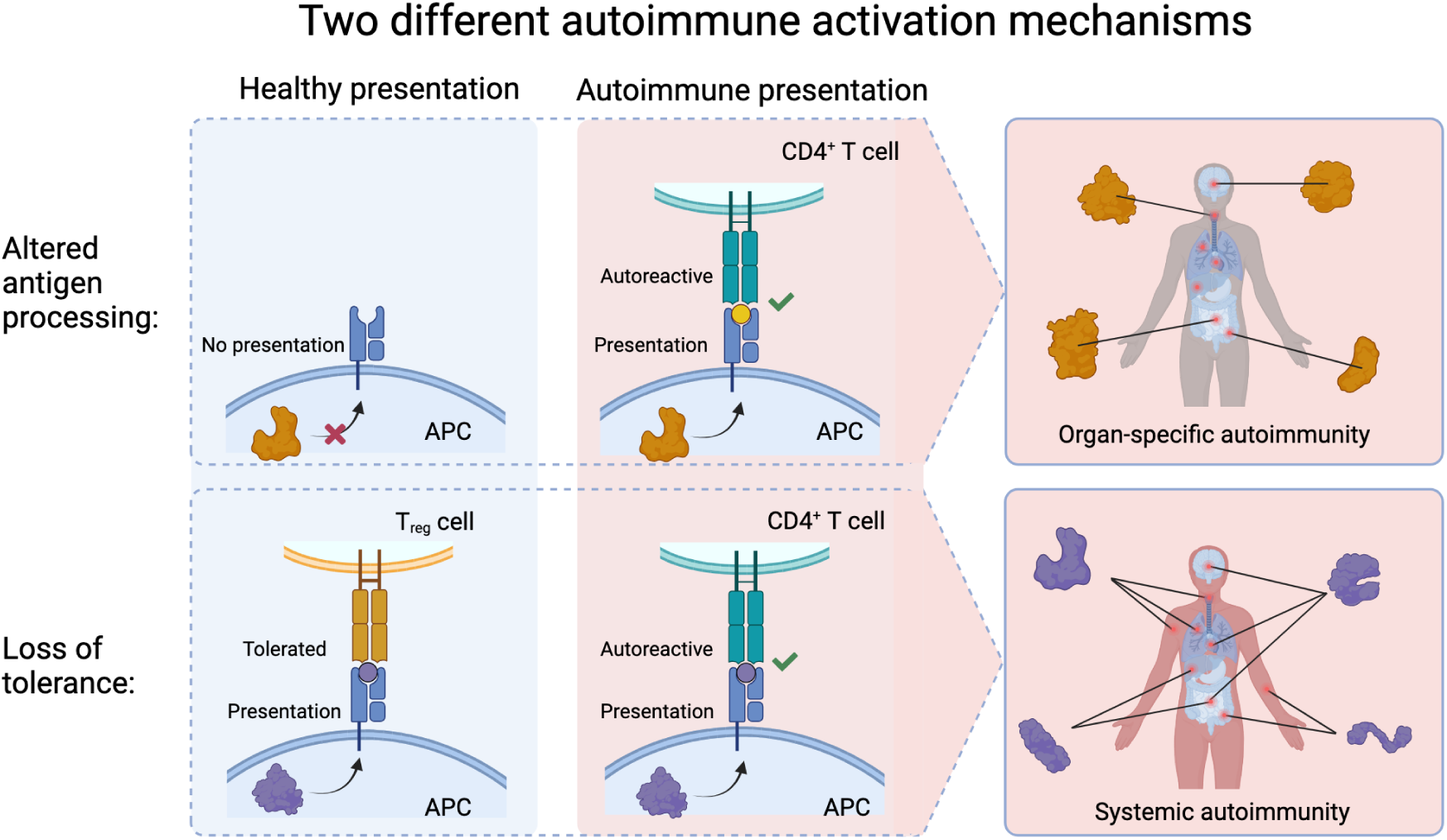

## Introduction

Autoimmune diseases arise when the immune system aberrantly targets self-proteins, resulting in organ dysfunction and tissue destruction (Song et al. 2024). Despite decades of research, the molecular events that trigger this loss of tolerance remain poorly defined. Although the immune system encounters thousands of self-proteins, only a limited subset become autoimmune targets, both in organ-specific diseases such as in myasthenia gravis (Vrolix et al. 2010) or multiple sclerosis (Kaushansky et al. 2010), and in systemic diseases such as lupus and rheumatoid arthritis (Wang et al. 2015, Ostrov 2015). This selectivity suggests that autoantigens possess intrinsic features that make them more likely to provoke autoreactive responses (Backes et al. 2011; D. Wang et al. 2017; Saksager et al. 2025). Understanding what distinguishes these proteins from the rest of the proteome may help identify early determinants of autoimmunity.

A defining feature of most autoimmune diseases is their association with particular HLA class II alleles. Genetic studies have consistently identified the HLA region to be the strongest susceptibility locus across autoimmune conditions such as type 1 diabetes, multiple sclerosis, rheumatoid arthritis, and systemic lupus erythematosus (Morran et al. 2015; Sawcer et al. 2014; Kochi et al. 2014; Langefeld et al. 2017). Because HLA class II molecules determine which peptides are presented to CD4⁺ T cells, these findings point to antigen presentation as a central checkpoint in disease initiation. However, a key question remains unresolved: does autoimmunity arise primarily from breakdown of tolerance toward normally presented self-peptides, or from altered antigen processing that reveals previously unseen “cryptic” peptides that escape tolerance induction? Under inflammatory or stress conditions, “cryptic” peptides could become processed and displayed, activating T-cells (Moudgil and Sercarz 2005). In addition, earlier studies suggested the role of post-translational modifications (PTMs) may alter the major histocompatibility complex (MHC)-exposed residues of autoantigens transforming the tolerance to self into autoreactivity (Lichti and Wan 2023; Kacen et al. 2023; Santambrogio 2023). Yet, the precise step in the MHC class II pathway that drives this association remains unknown.

Antigen processing within the MHC class II pathway is shaped by coordinated proteolysis, peptide loading, and intracellular trafficking. Dysregulation of proteases, HLA-DM/DO balance, or inflammatory conditions can alter which peptides are ultimately displayed. Although several studies have reported associations between components of the antigen-processing machinery and autoimmune disease, such as altered cathepsin expression in myasthenia gravis (Tolosa et al. 2003), or dysregulated HLA-DM/DO ratios in type 1 diabetes (Olsson et al. 2022), systematic studies of MHC-II peptide processing differences across diseases remain scarce. In addition, intrinsic protein properties including localization, structural stability, and physicochemical composition, may influence their likelihood of presentation (Kropshofer et al. 1996; Olsson et al. 2022; Pishesha et al. 2022; Saksager et al. 2025). Dysregulation of any of these components can shift which peptides are displayed (Nanaware et al. 2019; Alvaro-Benito et al. 2016, Pishesha et al. 2022).

In previous work we observed that peptides derived from potential autoantigens are more frequently presented on HLA-DR molecules in healthy donors than peptides derived from random human proteins. However, this enrichment was not uniform across all autoantigens and appeared to be driven by only a subset (Saksager et al. 2025). This heterogeneity suggests that autoantigens are not mechanistically homogeneous and may arise through distinct biological routes.

Historically, identifying disease-relevant T cell autoantigens has proven technically difficult. As a result, the majority of known autoantigens come from autoantibody research, and our understanding of the T cell antigen landscape remains incomplete (Prinz 2023). However, the advent of immunopeptidomics, mass spectrometry–based profiling of naturally presented peptides, has transformed this field (Ternette and Purcell 2018). This approach allows identification of peptides eluted directly from MHC presentation, revealing the repertoire of peptides available to T cells in health and disease. Immunopeptidomics thus provides a unique opportunity to explore whether autoimmune diseases differ not only in the self-proteins they target but also in the processing and presentation of those proteins.

To test whether distinct antigen-processing mechanisms contribute to autoimmunity, we classified experimentally verified T-cell autoantigens according to their natural presentation in healthy individuals and examined whether cryptic and tolerant autoantigens differ in molecular properties, cellular localization, and disease associations. We further analyzed immunopeptidomics datasets from rheumatoid arthritis and multiple sclerosis to determine whether autoimmune presentation is accompanied by systematic alterations in peptide processing signals.

## Materials and Methods

### Collection and processing of data

We collected autoimmunogenic T-cell epitopes from the The Immune Epitope Database (Vita et al. 2019) downloaded in June 2023. We selected autoimmune peptides that met the following criteria: associated with autoimmune diseases, positively tested in T cell assay on MCH class II (e.g. proliferation, binding to TCRs or cytokine release), tested in human hosts and where source proteins were from the human proteome. We then selected only the diseases which had strong or moderate levels of evidence of autoimmunity based on the autoimmune registry (Autoimmune Registry 2023). We removed peptides originating from T cell receptors and immunoglobulins. Finally we mapped the epitopes to the protein sequence of Uniprot ID provided, and removed the epitopes without exact match within the sequence. This resulted in 2,550 entries, distributed over 19 autoimmune diseases 116 autoantigens, and 1,191 unique epitopes.

We also collected the reviewed human reference proteome with 20,387 proteins from UniProt in April 2023 (UP000005640, (Breuza et al. 2016)). We removed proteins containing invalid amino acids, and proteins marked as putative (lacking experiential evidence), resulting in 17,618 proteins. We mapped the peptides back to the human proteome and nested the peptides. From the nesting we thereby had a measure of the MHC class II enrichment for each of the human proteomes. Further details are described in Saksager et al. 2025.

We collected the HLA-DR peptides of an in-house immunopeptidomics dataset of healthy volunteer donors. Monocytes isolated from peripheral blood mononuclear cells (PBMCs) of healthy donors were differentiated into immature dendritic cells, and matured overnight with LPS. HLA-DR–bound peptides were recovered by L243 immunoprecipitation, eluted, and sequenced by Liquid Chromatography-Tandem Mass Spectrometry (LC-MS/MS). Spectra were searched against the human reference proteome using an FDR of 1%, with each donor experiment replicated 2–12 times (for full protocol, see Attermann et al. 2021. To complement our in-house data, we queried the SysteMHC database (Huang et al. 2024) for immunopeptidomics datasets derived from autoimmune disease samples. Two suitable datasets were identified, corresponding to rheumatoid arthritis (Q. Wang et al. 2017) and multiple sclerosis (J. Wang et al. 2020). From both studies we collected the eluted peptides, and their identified source protein. We mapped each peptide back to the source protein sequence, if no exact match were found it was removed. We further discarded peptides with PTMs.

From IEDB we further collected human protein ligands of MHC class II from MS-LC/MS experiments conducted on healthy donors. We remapped the protein ids from the Ligand data and the in-house DC healthy donor data using R package ensembldb v. 2.22.0, EnsDb.Hsapiens.v79 (Rainer et al. 2019). We could then compare the protein IDs from the two datasets, and context of the peptides from different cell types. We combined cell types for B cells and epithelial cells across cell lines respectively. We only retained cell types with more than 1000 epitopes, and with a specific cell type annotated (removed tissue samples). We used UpSetR v. 1.14.0 to create the upsetplot. Finally, we extracted the context of the peptides from the uniprot sequence, after we had removed epitopes with post-translational modifications, undefined amino acids or Selenocysteine.

### Classification of cryptic and tolerant proteins on HLA-DRB

We classified the human proteome into two classes; cryptic and tolerant protein. If a protein had at least one nested peptide or singleton peptide presented on the HLA-DRB across all the healthy donors a protein was classified as tolerant, referring to its exposure towards the immune system without eliciting an autoimmune reaction. If a protein did not have any peptides (neither nested nor singletons) presented it was classified as cryptic, referring to its hidden state towards the immune system.

We then further categorised proteins found to be autoimmunogenic based on the T cell assays from IEDB as autoantigens, and the rest of the human proteome as innocuous (harmless) self.

### Protein feature collection

We used DeepLoc-2.0 to create a multilabel prediction of subcellular localisation of all proteins (Thumuluri et al. 2022). We included Nucleus, Cytoplasm, Extracellular, Mitochondrion, Cell membrane, Endoplasmatic reticulum, Golgi apparatus, Lysosome/Vacuole and Peroxisome, but removed chloroplast/plastid as this is not found in human cells. The location with the highest likelihood was chosen as the primary subcellular localization for each protein, no proteins had peroxisome as the most likely location.

For the secondary structure and relative accessible surface (RSA), we used NetSurfP-3.0 and AlphaFold2 predictions (Høie et al. 2022; Jumper et al. 2021). NetSurfP-3.0 uses the protein sequence and returns the eight state secondary structure and RSA for each amino acid in each protein. We obtained the predicted Alphafold structures for the proteins by downloading PDB files corresponding to the same reference proteome from the AlphaFold predictions database. For each PDB file, we then used Biopython (Cock et al. 2009) to extract structure information from each PDB. From both NetSurfP and AlphaFold we converted the output by extracting the DSSP codes for the secondary structure and RSA for each amino acid.

We then created a sliding window of 7 amino acids, in which we counted the occurrence of each secondary structure. We counted each structure as truly present, when the frequency were above 5 for alpha-helix, 3 amino acids for β sheet, 3 amino acids for 3_10_ helix, 6 amino acids for π helix, 1 amino acid for turn, 1 amino acid for bend and 1 amino acid for coil (Schulz and Schirmer 1979). The ends were extended to prevent negative bias of structures placed there.

We collected and calculated the proteins’ physico-chemical properties through two different packages. Firstly, we used Biopython to calculate molecular weight, protein length, protein aromaticity, protein instability index, hydropathic character of the proteins with the KyteDoolitttle scale, the grand average hydropathy which is the hydropathicity normalised by protein length, and the isoelectric point and charge at ph 5 and 7 to mimic the environment in the cell cytosol and lysosome (Cock et al. 2009; Kyte and Doolittle 1982). Lastly, we computed the molar extinction coefficient in both the reduced and oxidized form. Aromaticity is the relative frequency of aromatic amino acids (Phe, Tyr and Trp) based on Lobry and Gautier 1994, while the instability index was based on the work by Guruprasad et al. 1990. The instability index provides an estimate of protein stability by summing the weighted contributions of all dipeptides in the sequence, normalized by protein length.

Secondly, we obtained protein features based on the R packages Peptides v. 2.4.6 (Osorio et al. 2015). Including protein length, molecular weight, isoelectric point, hydrophobicity, aliphatic index, instability index, and hydrophobic moment. Further we calculated the number of amino acids and the molecular percentage of the amino acids groups: Tiny [A,C,G,S,T], Small [A,C,D,G,N,P,S,T,V], Aliphatic [A,I,L,V], Aromatic [F,H,W,Y], NonPolar [A,C,F,G,I,L,M,P,V,W,Y], Polar [D,E,H,K,N,Q,R,S,T], Charged [D,E,H,K,R], Basic [H,K,R] and Acidic [D,E]. Lastly, we calculated the composition of amino acids as a percentage of protein length.

We classified the investigated autoimmune diseases into organ-specific or systemic categories based on a targeted literature review. First, we identified two key papers that explicitly discussed autoimmune diseases in each category Ostrov 2015 and Wang et al. 2015. The papers served as primary references for classification. For each disease, we then searched for at least one additional publication that supported its categorization. Diseases not covered in either of the primary references were classified based on at least two independent sources that consistently assigned them to the same category. In cases where we encountered publications that contradicted the established classification, these were noted separately as opposing evidence.

### Peptides secondary structure and relative solvent accessibility

The peptides from LC-MS/MS of healthy donors, rheumatoid arthritis and multiple sclerosis patients were mapped onto their corresponding source protein sequences, and per-residue relative solvent accessibility (RSA) and secondary structure (SS) assignments were retrieved from pre-computed protein-level data (described previously). For each peptide, the RSA values spanning its sequence positions were extracted, and the median RSA was calculated to represent the peptide’s overall solvent exposure. For each peptide, the proportions of secondary structure types were computed by counting occurrences of each symbol in the SS annotation and dividing by peptide length. Comparisons of median RSA and SS composition fractions between disease and healthy groups were performed using two-sided Mann–Whitney U tests. P-values were adjusted for multiple testing with the Benjamini–Hochberg false discovery rate (FDR) procedure, and Cliff’s delta was calculated to estimate effect sizes.

### Statistics

Categorical variables, specifically the counts of proteins across subcellular locations and disease phenotypes, were compared between classes using Chi-square tests of independence. For subcellular location, the expected background distribution was defined as the overall distribution of proteins across all locations in the complete dataset. For disease phenotype, an equal distribution across the different disease categories was assumed as the background. In the cases where the same autoantigen has been investigated in multiple autoimmunities, it was counted in all diseases.

Significance was further evaluated by examining the adjusted standardized residuals of the Chi-square test, as defined by Agresti 2008. According to this definition, residuals around ±2 can be considered significant at the 5% level in tables with relatively few cells, whereas in larger tables, a more conservative threshold of around ±3 is recommended. Given that our analysis involved 34 cells, we considered this to fall into the “many cells” case and applied the ±3 criterion when interpreting the results.

Continuous structural features, including physicochemical and geometric descriptors, were analyzed across the four protein classes. Since the features were not normally distributed, non-parametric Kruskal–Wallis tests were applied. Where significant differences were detected, post hoc pairwise comparisons with Holm-Bonferroni multiple-testing correction were performed to identify which classes differed. Effect sizes were additionally calculated with a dunn-test to provide an estimate of the magnitude of observed differences, complementing significance testing.

For the physico-chemical properties, we compared the median values of these features across groups, to statistically assess differences between classes. For amino acid composition, CLR-transformed data were analyzed using analysis of variance (ANOVA), while physico-chemical properties were tested using the non-parametric Kruskal–Wallis test. Where global tests indicated significant differences, we performed post-hoc pairwise analyses: Tukey’s test following ANOVA, and Dunn’s test following Kruskal–Wallis, to identify the specific groups contributing to the observed effects.

### GO terms and network analysis

Based on the classification of the autoantigens into cryptic and tolerant proteins, we analysed the GO term enrichment of each of the two groups. We used g:Profiler’s g:GOSt functional profiler with the default settings (Kolberg et al. 2023), using Homo sapiens as organism, only annotated genes and a significance threshold 0.05 based on g:SCS. We downloaded the GO Biological process (BP) results on the 7th of August 2025.

Using the R package OntologyIndex v. 2.12 (Greene et al. 2017), we connected the direct Parent:Child terms of the enriched terms. Based on the g:Profiler, we had the connection of each gene to the BP, and the genes were associated with autoimmune diseases from the experimental data from IEDB. The two networks of the tolerant and cryptic autoantigens were then visualised using Cytoscape (Shannon et al. 2003). The BP Parent:Child edges were colored with green directed arrows, while each gene associated with the enriched term were illustrated with a grey undirected edge. The nodes were colored according to the associated disease, in case of multiple diseases it was depicted as a pie-chart. The genes and diseases which were not associated with an enriched term were not included in the illustration.

### Sequence logo generation and comparison

We had previously mapped each peptide from the healthy donors, rheumatoid arthritis and multiple sclerosis to their source protein. We then extracted the context from the peptides, three amino acids up- and down-stream of the C- and N-terminal sites of the peptide. The peptide contexts of each dataset were used as input to Seq2Logo 2.0 (Thomsen and Nielsen 2012). It was run as p-weighted Kullback-Leibler plot, no clustering and a 200 weight on prior. We additionally to the plot, also extracted the PSSM and p-weighted KL matrix of each dataset. Based on the p-weighted KL matrices we calculated the Kullback-Leibler divergence (KLD) per position of the context, 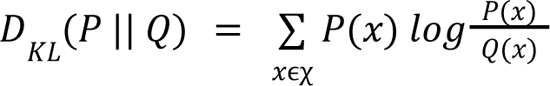, where the healthy donor peptide distribution is Q and each of the disease distributions is P. The KL-divergence can be interpreted as a measure of how likely it is that the distribution P can come from Q. We also sampled the 1400 peptides from the healthy donors 1000 times with replacement from the healthy donors, extracted the p-weighted KL matrices from each of these sets and calculated the KLD for each. For each position of the context we then used it as a null-distribution on which we calculated an empirical non-parametric p-test and the lower and upper bound of the 95% confidence intervals.

Additionally, we ran IceLogo (Colaert et al. 2009) with each of the disease dataset as the experimental data, and the healthy donor peptides as user-submitted reference data using percent difference and a significance threshold of p < 0.05.

We conducted the same KL analysis for the context of the LC-MS/MS ligand data from IEDB for the cell types: B cells, myeloid cells, and epithelial cells.

## Results

### Classification of proteins based on presentation and autoimmunogenicity

We divided the entire human proteome into two categories: proteins with at least one peptide presented in a healthy donor, and proteins which are not found to be presented in any of the healthy donors. If a protein was presented it is denoted as *tolerant*, as it is presented in healthy conditions and must therefore be tolerated by the immune system. The proteins which were not presented we denote as *cryptic*, these proteins cannot trigger an immune response since they are hidden from the immune cells. We remark that the healthy donor immunopeptidomics dataset contains a high number of replicates of the same experiment, ensuring high coverage of all presented peptides.

Additionally, we subcategorised proteins which were described as autoantigens, according to the set of positive T cell assays from IEDB, we had enrichment data for 103 autoantigens. If a protein has been shown to have at least one autoimmunegenic T cell epitope it is denoted as an Autoantigen. The remaining human proteome is denoted as Innocuous Self. The final division of the human proteome into these four classes is illustrated in Fig. 1.

**Fig. 1.**
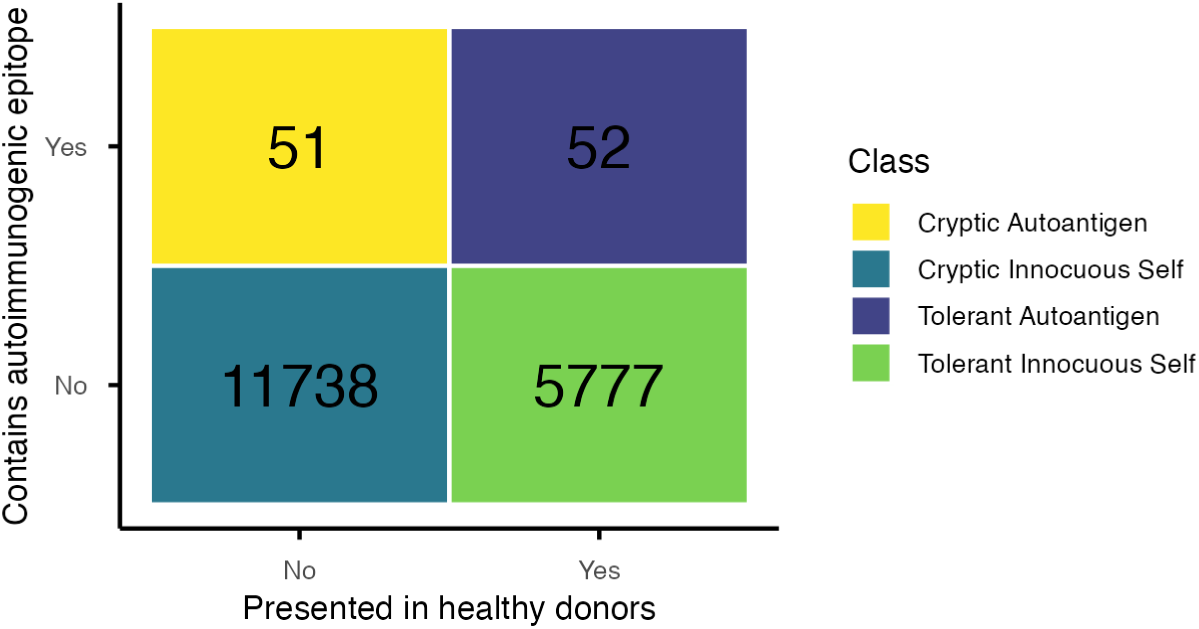
Classification of the human proteome. The x-axis divides the proteins into proteins which have at least one peptide presented on dendritic cells in across 40 healthy donors. The y-axis divides the proteins into proteins which have at least one epitope with autoimmunogenic properties based on T cell assays of autoimmune patients.

The classification results in four unbalanced groups, showing that two-thirds of the human proteins are not presented, while one-third of the proteome is presented on HLA-DR in healthy donors. This distribution is not repeated between the autoantigens. The size of the two groups cryptic and tolerant autoantigens are fairly equal.

Because the autoantigens are represented in both classes we hypothesise that there are two mechanisms by which proteins are turned into autoantigens. The first are cryptic proteins, which under normal conditions are not presented on MHC to the immune system. Consequently, T cells remain untolerized and unactivated under normal conditions. Due to the fact that autoimmune responses have been found to these autoantigens, it is a natural interpretation that in autoimmunity these cryptic autoantigens are, for unknown reasons, presented in autoimmune patients.

The second mechanism is for the tolerant autoantigens, which under normal conditions are presented but do not elicit an autoimmunogenic T cell response. Here a response can be caused by a different set of presented peptides from the self protein or due to the breakup of the CD4^+^ T cells tolerance response. We hypothesize that the Cryptic autoantigens and the Tolerant Autoantigens arise from two inherently different mechanisms. In order to understand which properties shape these different autoantigens, we have analysed cellular localisation, physico-chemical properties, and performed a network analysis of the two groups of autoantigens to better characterise their different origins.

### Cryptic autoantigens are found in the cell membrane

Self-proteins subcellular localisations can influence their uptake into the MHC class II presentation pathway due to the different uptake mechanisms. We previously found that autoantigens were often placed in cell membranes and extracellular spaces (Saksager et al. 2025).

We have here examined potential differences in the subcellular localization of cryptic and tolerant autoantigens. The most probable localization of each human protein was predicted using DeepLoc-2.0, which provides reliable estimates of proteins’ actual cellular compartments. We then assessed whether the four protein groups (as defined in Fig. 1) differed significantly in their localization patterns. A comparison between cryptic and tolerant autoantigens revealed striking contrasts. Cryptic autoantigens were markedly enriched in cell membranes and lysosomal or vacuolar compartments, with a corresponding depletion in nuclear proteins. In contrast, tolerant autoantigens were predominantly localized to extracellular regions. Specifically, we used the entire human proteome as a background distribution of subcellular localisations, and then tested the actual distribution of each category against the expected with a chi-square test (Fig. 2). We considered the standard residuals significant at a 5% level at a threshold of ±3 in concordance with the definition by Agresti (Agresti 2008).

**Fig. 2.**
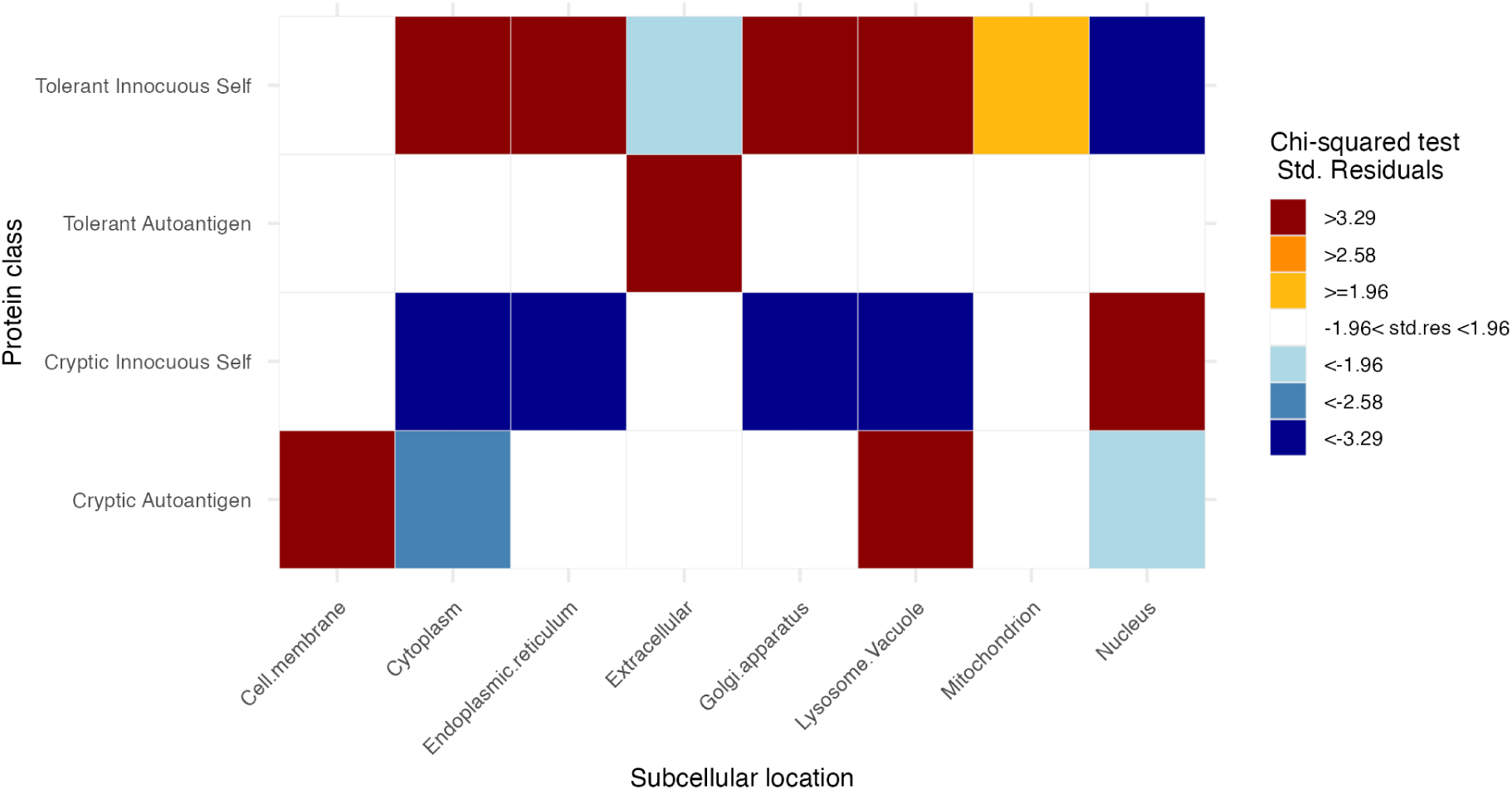
The difference of subcellular localisations for cryptic and tolerant autoantigens and innocuous self proteins. The standard residuals of the chi-square-test between the four classifications of human proteins across their primary subcellular localisation predicted by DeepLoc-2.0. A standard residual above |3.29| is considered significant for many variables according to Agresti, 2008 (Agresti 2008).

Fig. 2, also shows that two groups, tolerant and cryptic innocuous self, almost reciprocate to each other. The tolerant innocuous selves are mostly found in the cytoplasm, endoplasmic reticulum, golgi apparatus and lysosomes/vacuoles, while the cryptic innocuous selves are strongly underrepresented in these sites. Further the proteins found in the nucleus are highly overrepresented in the class which is not presented on healthy donors, in agreement with previous research (Schmid and Münz 2005).

The enrichment of cryptic autoantigens within the lysosomal/vacuolar category should, however, be interpreted with caution, as DeepLoc-2.0 predictions of this location have the lowest prediction performance (Thumuluri et al. 2022). It is likely that some membrane-associated proteins are predicted as lysosomal proteins. Lastly, the two types of cryptic, autoantigens and innocuous self, as well as the two tolerant types are also distinguishable in their locations. This means that both cryptic and tolerant autoantigens potentially can be separately identified from the remaining human proteome.

The clear distinction between membrane-bound cryptic autoantigens and extracellular tolerant ones suggests that intrinsic protein properties influence their likelihood of presentation. These differences may reflect distinct uptake and processing routes, linking subcellular localization to both protein function and predisposition for MHC class II presentation and autoantigenicity.

### Cryptic and tolerant innocuous self proteins have differences in physico-chemical properties

To determine whether intrinsic protein features affect MHC class II presentation, we compared autoantigens and other self-proteins across secondary structure, amino acid composition, and physicochemical properties that influence proteolytic processing and presentation efficiency. We retained the division of proteins into the four predefined classes, as outlined in Fig. 1. To characterize differences between these groups, we quantified a broad set of sequence-derived features, listed in Methods.

All physicochemical features tested differed significantly between the presented and non-presented innocuous self-proteins (p < 0.05), showing that intrinsic molecular properties strongly influence whether proteins are displayed by MHC class II. This confirms that antigen presentation is not random but shaped by biochemical constraints such as amino acid composition, charge, solubility, and structural stability, which likely are affecting susceptibility to proteolysis and trafficking into endosomal compartments.

When comparing autoantigens to their respective innocuous counterparts, however, the differences were minimal. Among cryptic proteins, only glutamine content was significantly lower in autoantigens (p < 0.05), though the effect size was small. For tolerant proteins, autoantigens contained slightly more glycine and arginine, fewer leucine residues, and showed an enrichment of tiny amino acids (A, C, G, S, T) alongside a depletion of aromatic residues (F, H, W, Y). These patterns suggest that tolerant autoantigens may be somewhat more flexible and hydrophilic, but the magnitude of change is limited.

The differences between autoantigens and innocuous self-proteins were generally few and small, with limited explanatory power for why certain proteins become immunogenic (Supplementary Material 1, S.Fig1). The molecular extinction coefficients were higher for cryptic autoantigens than for innocuous self-proteins and lower for tolerant autoantigens. Aromatic amino acids such as tyrosine and tryptophan have high extinction coefficients (Pawar and Bichile 2011), this therefore matches the significant depletion of aromatic residues in the tolerant autoantigens. The variation in molecular extinction coefficients could therefore represent differences in their structural stability and degradation dynamics during antigen processing.

In summary, while physicochemical features clearly define which proteins are likely to be presented by MHC II, they do not distinguish autoantigens from other self-proteins. These results imply that additional factors, such as subcellular localization, folding dynamics, or antigen-processing differences, play a more decisive role in determining autoantigenicity.

### Tolerant autoantigens are highly connected to immune processes

The clear differences in subcellular localization suggest that the proteins may participate in distinct cellular functions. In general, localization and function are closely linked in biology, as the compartment in which a protein resides determines its molecular interactions, processing environment, and biological role.

We therefore analysed the enrichment of gene ontology terms connected to the two sets of autoantigens, cryptic and tolerant. For each set of autoantigens we made a GO term enrichment analysis, and selected all biological processes with a significance-threshold of 0.05. Using OntologyIndex we inferred which of the significantly enriched GO terms had a parent:child relationship (Greene et al. 2017). The autoantigens were further connected to autoimmune diseases through the experimental data from IEDB, as previously described. The network connecting the proteins to the GO terms, and the relationship between the GO terms are visualised in Fig. 3.

**Fig. 3.**
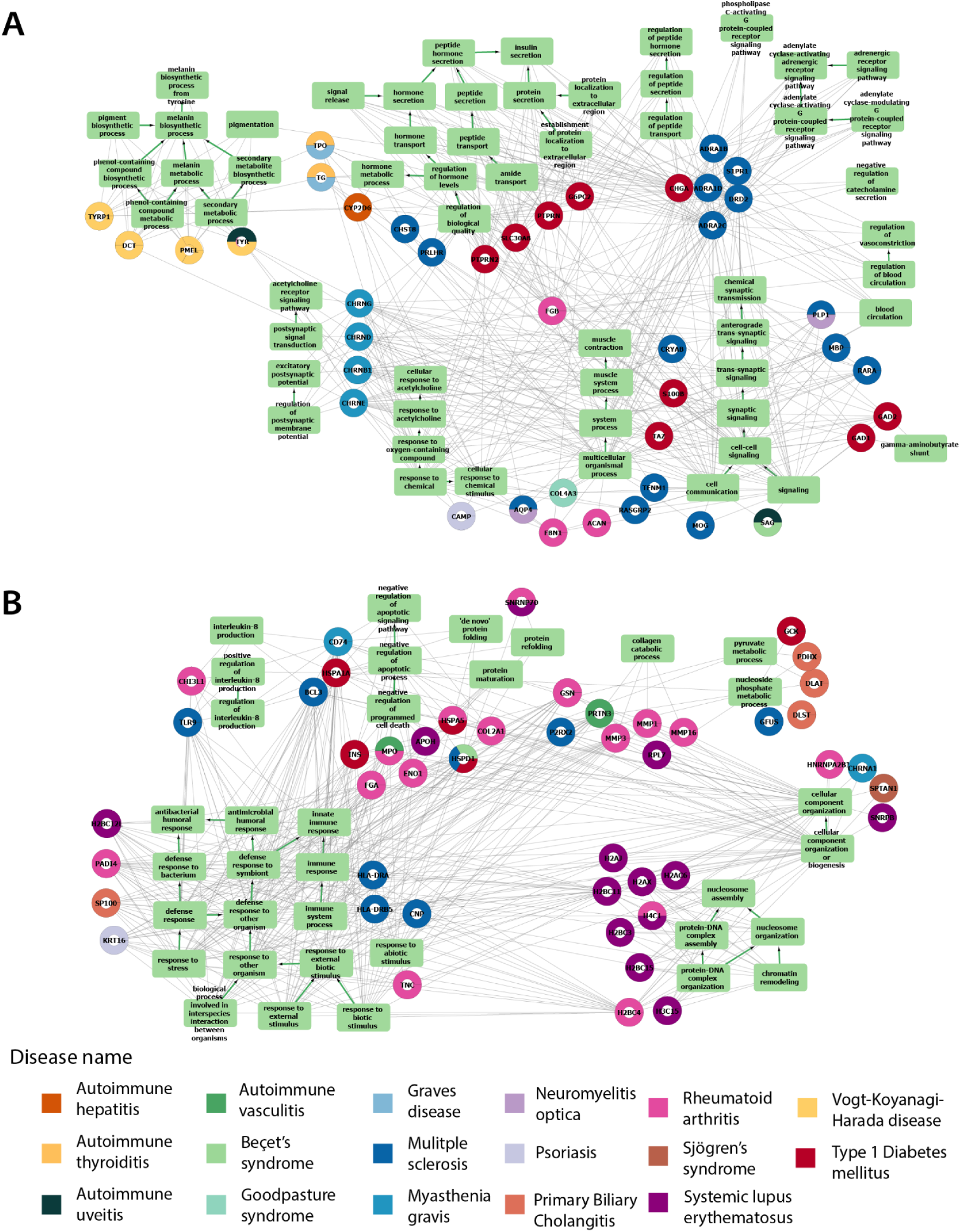
Networks of significantly enriched Gene Ontology biological processes of autoantigens. The square nodes are enriched biological functions, connected with each other in parent-child relations indicated with green arrows as edges. The round nodes are the autoantigens connected with grey edges to the GO terms. Each autoantigen is colored by associated autoimmune disease, and if the autoantigen has been investigated in multiple diseases the colors are split. Note, if an autoantigen is not connected to at least one enriched term it is not shown in the network. (A) Cryptic autoantigens and their associated biological processes (B) Tolerant autoantigens and their associated biological processes.

The results showed clear differences between the tolerant and cryptic autoantigens (Fig. 3). The cryptic autoantigens are dominated by T1D and MS (Fig. 3A), while Lupus and RA are prominent for the tolerant autoantigens (Fig. 3B).

The tolerant autoantigens show a clear pattern that almost solely contains biological processes associated with immune responses and cell death; *immune response*, *defence response to other organisms*, *response to stress*, *response to external biotic stimulus*, and n*egative regulation of cell death* and *negative regulation of apoptotic signalling pathway*. The immune response terms are highly connected to thirty autoantigens across nine diseases, while the apoptotic terms are eleven autoantigens from seven different diseases. Several proteins across diseases are also associated with cytokine production, in particular *interleukin 8.* The few other enriched terms are associated with protein folding and cellular organisation, but are not as widely connected to the autoantigens.

The cryptic autoantigens do not show the same extent of association with immune responses. Instead they appear more disease specific and related to the targeted and degraded tissues. There are for instance terms such as *insulin secretion,* and *adrenergic receptors signalling pathway*, which are highly related to the disease mechanism of type 1 diabetes mellitus in which the beta-cells which produce the hormone insulin are attacked (Ilonen et al. 2019).

The term *Melanin biosynthetic process* is associated with all the Vogt-Koyanagi-Harada genes, a disease which targets melanocytes which produce pigment (Sakata et al. 2014). There is also a clear cluster of myasthenia gravis genes related to the *acetylcholine receptor signaling pathway*, and *synaptic transmission* (Fig. 3A). Myasthenia gravis is characterised by autoantibodies binding to the acetylcholine receptor and thereby interfering with the signalling (Tannemaat et al. 2024; Vrolix et al. 2010).

For genes related to autoimmune hepatitis, autoimmune thyroiditis and Grave’s disease there are terms such as *hormone metabolic process* and *regulation of hormone levels*. Autoimmune hepatitis, as the name indicate, targets the liver cells which functions include removal of excess hormones, thyroiditis degrades the thyroid gland which produces the hormones T_3_ and T_4_, and in Grave’s autoantibodies binds to the receptor of the thyroid resulting in over-expression of hormones.

We note that the GO enrichment analysis is likely skewed by diseases with many known autoantigens (e.g. MS, RA, T1D), which dominate the annotation space and can mask subtler trends in less-studied diseases.

In the tolerant autoantigen network, there is a single disease-specific cluster around *nucleosome assembly*, which seems mostly associated with systemic lupus erythematosus. Lupus is in general known to target histones (Pieterse and Van Der Vlag 2014). The complete results of the GO term analysis is provided in Supplementary Material 2 and 3.

The contrast between the cryptic and tolerant autoantigens’ enriched GO terms, reveals a fundamental biological distinction between the two types of proteins. While the tolerant autoantigens align with immune system regulation and cell death control, the cryptic autoantigens reflect processes intrinsic to the targeted tissues, consistent with disease-specific pathology. We note that the Gene Ontology enrichment analysis highlights the dominant biological themes but may underrepresent functions associated with diseases from which we have few autoantigens.

The distinct biological processes and cellular localizations suggest that the proteins’ physiological roles shape their exposure to antigen-processing pathways. Proteins central to immune regulation may be constitutively processed and presented, whereas tissue-specific proteins could become antigenic only under altered processing or stress conditions.

### Cryptic and tolerant autoantigens divides autoimmune phenotypes

The network visualisation (Fig. 3) indicated a potential link between disease type and autoantigen class. Notably, rheumatoid arthritis and lupus were dominating among tolerant autoantigens, while multiple sclerosis and type 1 diabetes seemed more prevalent within the cryptic group. To further understand this pattern, we next investigated how autoantigen distribution relates to the traditional classification of autoimmune diseases into organ-specific and systemic forms.

Based on how each autoimmune disease was commonly described in the literature, we divided the diseases into two groups, organ-specific and systemic (Supplementary Material 1 S.Table 1). We then sorted the autoantigens based on their association with the autoimmune diseases, Fig. 4A. We have twelve organ-specific diseases with a total of 65 autoantigens, 42 cryptic and 23 tolerant, and seven systemic diseases with 41 autoantigens divided between 10 cryptic and 31 tolerant autoantigens. All seven systemic diseases have more tolerant autoantigens than cryptic ones, except Vogt-Koyanagi-Harada disease which do not have any tolerant autoantigens. The opposite is seen for the organ-specific diseases, where nine out of the ten diseases have more cryptic autoantigens than tolerant autoantigens. The two exceptions are primary biliary cholangitis and autoimmune hepatitis, where the latter have an equal amount in each class.

**Fig. 4:**
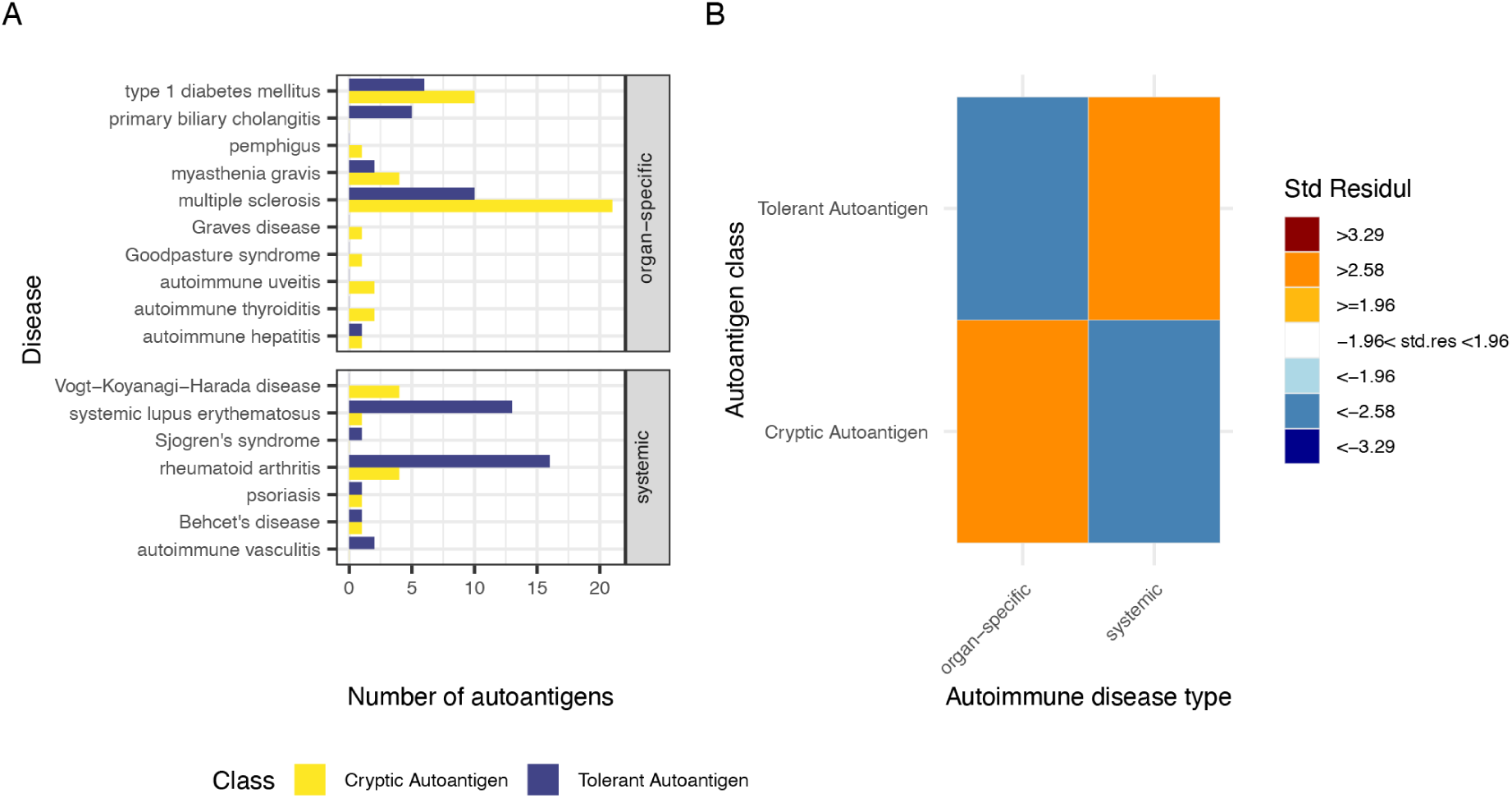
(A) The number of autoantigens for each of the autoimmune diseases distributed by disease phenotype, organ-specific and systemic disease. (B) Standard residuals of a chi-square test between disease phenotype and associated autoantigen classes. A standard residual above |2.58| is considered significant for few variables (Agresti 2008).

**Table 1:** Overview of the three immunopeptidomics datasets.

| Disease | Disease Type | Number of Patients | HLA subtypes | Method | Tissue/Cell types | Total number of peptides |
| --- | --- | --- | --- | --- | --- | --- |
| Rheumatoid Arthritis | Systemic | 5 | DRB1: 0101,0401, 0402,0801, 1104, 1501 | LC-MS/MS: LTQ-Orbitrap XL, Q exactive plus, 6550 iFunnel QTOF | Synovial tissue and fluid, and PBMCs | 578 |
| Multiple Sclerosis | Organ-specific | 3 | DRB1*15:01<br>DRB5*01:01 | LS-MS/MS: LTQ Orbit XL | B cells and macrophages | 1,829 |
| Healthy donors | Healthy | 40 | DRB1, DRB3, DRB4, DRB5 | LC-MS/MS: QExactive HF orbitrap | Dendritic cells | ~ 80,000 |

We used a chi-square test, to examine if the autoantigens classification of tolerant/cryptic is independent of the autoantigens disease category. The standard residuals are shown in Fig. 4B.

A value ± 2.58 represents a significant difference at the 5% level, since there is a small number of groups in the test (Agresti 2008). All four residuals are above that threshold: Cryptic autoantigens are highly overrepresented in the organ-specific diseases, while there are more tolerant autoantigens in the systemic group. This finding, that the phenotype of an autoimmune disease can be related back to whether an autoantigen is presented under normal conditions, supports the model of a meaningful distinction between cryptic and tolerant autoantigens. It further suggests that there might be two different mechanisms off-setting autoimmunity and that this results in either organ-specific or systemic diseases.

It is worth noting that although the classification reveals a statistically significant division between the diseases, it does not represent an absolute separation. Most of the diseases have autoantigens in both classes, including T1D, Lupus, RA, psoriasis, MG, MS, and autoimmune hepatitis. Nevertheless, several diseases with a low number of associated autoantigens, have autoantigens from only a single class. For instance, autoimmune uveitis, and autoimmune thyroiditis have only cryptic autoantigens, while autoimmune vasculitis and Sjögrens’ have only tolerant autoantigens. Due to the low number of autoantigens in each individual disease we could not meaningfully test the distribution for the individual diseases.

The distinction between organ-specific and systemic autoimmune diseases cannot be applied systematically, as the definition is diffuse and based largely on clinical presentation rather than underlying mechanisms. Furthermore, patients may present with heterogeneous autoantigen profiles, complicating strict classification. In our analysis, we also included psoriasis, a condition debated as autoimmune versus autoinflammatory; given its frequent description as an immune-related systemic disease, we classified it as systemic. We also decided to exclude Neuromyelitis optica’s two cryptic autoantigens, due to lack of description as either organ-specific or systemic in the literature, combined with the recent conflicting findings of (Samim et al. 2024), where ∼31% of the patients showed constitutional symptoms.

### Immunopeptidomics studies of Rheumatoid Arthritis and Multiple Sclerosis

To determine whether differences in antigen processing contribute to the emergence of autoimmune epitopes, we next analyzed immunopeptidomics datasets from two autoimmune diseases. While previous analyses focused on the intrinsic features of autoantigens, immunopeptidomics allows direct examination of the peptides that are naturally processed and presented on MHC class II molecules. This enables us to assess whether disease-specific alterations in processing or presentation can be observed in vivo.

We analyzed two available immunopeptidomics datasets from autoimmune diseases. One dataset was derived from the organ-specific disease multiple sclerosis (J. Wang et al. 2020), which primarily targets the myelin sheaths of the central nervous system. The other was from rheumatoid arthritis (Q. Wang et al. 2017), a systemic autoimmune disease characterized by inflammation of the synovial joints with frequent extra-articular manifestations. These studies aimed to identity HLADR-presented self-peptides in cells taken from clinical samples by isolating the MHC:peptide complex and subsequently running a liquid chromatography-tandom mass spectrometry (LC-MS/MS) (J. Wang et al. 2020; Q. Wang et al. 2017), comparable to the healthy donor data (Attermann et al. 2021).

There are, however, differences in sample size, cell types analyzed, HLA repertoires, sequencing depth, and number of replicates, Table 1. Due to the limited number of immunopeptidomics datasets on autoimmune diseases available it was not possible to find any datasets with more similar experimental setups. Consequently, our analysis focuses on peptides that are identified in the disease datasets rather than on the absence of peptides relative to healthy controls. In contrast, the healthy dataset is considered broadly representative of the normal immunopeptidome, owing to its large cohort size, broad HLA coverage, and multiple technical replicates. The healthy donor dataset includes 40 distinct HLA-DRB alleles, encompassing both DRB alleles observed in the multiple sclerosis samples and all six DRB alleles from the rheumatoid arthritis patients. Thus, peptides identified in the patient datasets are unlikely to arise solely from differences in HLA binding affinity relative to the healthy controls. Lastly, we address in more detail the differences between presentation of peptides within cell types in later in this paper.

### Different antigen processing signals in peptides presented in autoimmune disease

Given our hypothesis that altered antigen processing contributes to autoimmune presentation, we examined the sequence context surrounding peptide termini in rheumatoid arthritis (RA) and multiple sclerosis (MS) relative to healthy donors. Peptide termini correspond to proteolytic cut-sites, and their local amino acid composition can reveal changes in antigen processing or MHC loading preferences. For each dataset, all identified epitopes were mapped back to their source proteins. We extracted the three amino acids upstream of the peptide N-terminus and downstream of the C-terminus, along with the first and last three residues of each peptide. Sequence preferences were visualized using position-weighted Kullback–Leibler (KL) logos, Fig. 5A-C.

**Fig. 5.**
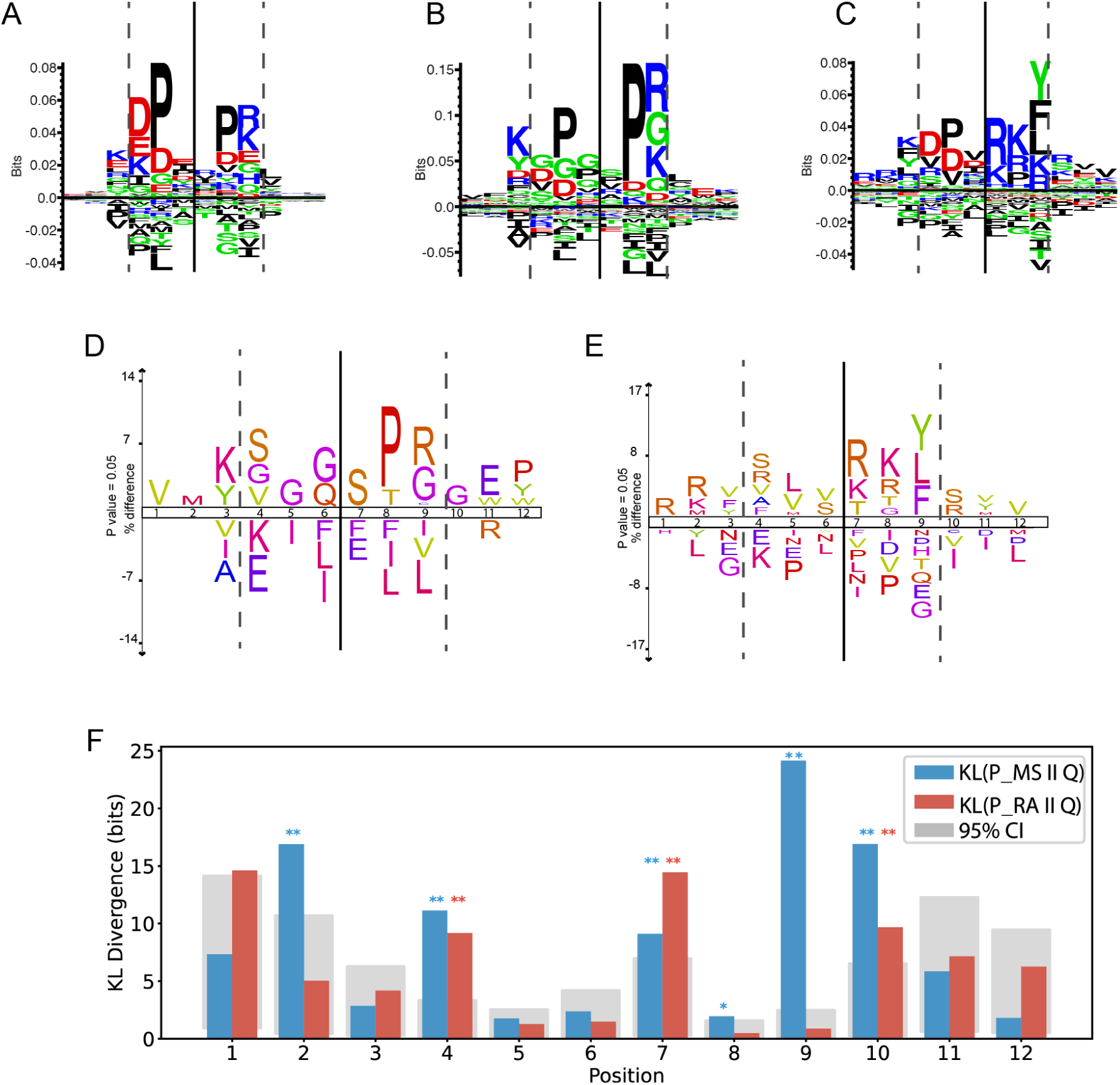
Comparison of the peptide sequence context between peptides presented in healthy donors, rheumatoid arthritis patients and multiple sclerosis patients. Position 1-3 is the upstream and position 4-6 downstream amino acids from the peptides N-terminal site, while the position 7-9 is upstream and 10-12 is downstream of the C-terminal site. Probability-weighted Kullback-Leibler sequence logo plot of healthy donors (A), rheumatoid arthritis (B) and multiple sclerosis (C) created with Seq2Logo. ICElogos with the disease percent-differences to the healthy, RA (D) and MS (E). (F) The barplot shows the measured KL divergence between the probability-weighted Kullback-Leibler matrices. The red bars are the KL divergence from the healthy donors to the rheumatoid arthritis, and the blue is the divergence from the healthy to the multiple sclerosis. We further estimated the confidence intervals of the KLD by sampling a 1000 random sets from the healthy and measuring their KLD to the healthy. The 95% confidence interval is shown as the grey area for each position. Significant difference at p > 0.05 is marked by *, and p > 0.01 is marked by **.

In healthy donors, sequence preferences closely matched previously reported antigen processing motifs (Egholm Bruun Jensen et al. 2024), with strong enrichment for acidic (D, E) and basic (K) residues at the N-terminal position, and a prominent proline at the second position. The C-terminal region showed enrichment for P and D at the 8th and 9th position in the plot, followed by positively charged residues (R, K) at the peptide terminus. RA peptides largely maintained this canonical pattern, though with subtle shifts, an increased frequency of glycine (G) at several N-terminal positions and enhanced prominence of proline at the C-terminal penultimate site. MS peptides, in contrast, exhibited broader deviations: weakened proline preference at both termini, an enrichment of R and K across the C-terminal region, and a distinct emergence of hydrophobic residues (Y, F, L) at the terminal position.

To assess whether these differences were statistically significant, we measured the Kullback-Leibler divergence (KLD) from the healthy donors’ peptide context (p-weighted KL) to each of the disease contexts per position, Fig. 5F. The KLD is a measure of how different one distribution is from another. We then drew 1000 random sets of peptides from the healthy donors and calculated their KLD to the healthy a get a null-distribution of KLD per position. The 95% of the confidence intervals are smallest at the positions with the most information, eg. position 4, 5, 8 and 9, representing that if we draw a random sample the distance from the background to the sample is small. We see a significant difference for MS at position 2, 4, 7, 8, 9 and 10, and a significant difference at 4, 7 and 10 for rheumatoid arthritis. This suggests that the processing signal is different in autoimmune disease, especially for multiple sclerosis.

Additionally, we generated two ICElogos (Colaert et al. 2009) using the healthy donor dataset as a reference and performed z-tests at a 5% significance level. All differences described above were statistically significant (Fig. 5DE). The magnitude of change reached up to 14% per site in RA and 17% per site in MS. While RA displayed fewer positional deviations overall, individual changes were more pronounced compared to MS. The PSSM logos of each of the three sets can be seen in Supplementary Material 1 S.Fig2).

Overall, these results suggest that while RA maintains largely physiological antigen-processing patterns, MS shows evidence of altered proteolytic trimming and peptide generation.

### Antigen processing signals does not vary between different cell types

Before comparing peptide processing patterns in autoimmune disease, we first evaluated whether the observed differences could reflect variation in the antigen-presenting cells (APCs) rather than disease-specific effects. Because the datasets originated from distinct sources—dendritic cells (DCs) from healthy donors and synovial tissue or PBMC-derived APCs from patients—it was essential to confirm that the cell type itself did not bias the antigen-processing signatures. Variations in the proteolytic machinery, such as the expression of different cathepsins, could theoretically alter the peptide-flanking sequences visible in the MS data (Fig. 5).

To determine whether our healthy donor DC dataset provided representative coverage of the human immunopeptidome, we compared it with all available MHC class II eluted ligand datasets from the Immune Epitope Database (IEDB), including studies from B cells, macrophages, dendritic cells, splenocytes, and thymocytes. We visualized the overlap between the proteins identified in these cell types using an Upset plot (Fig. 6A).

**Fig. 6.**
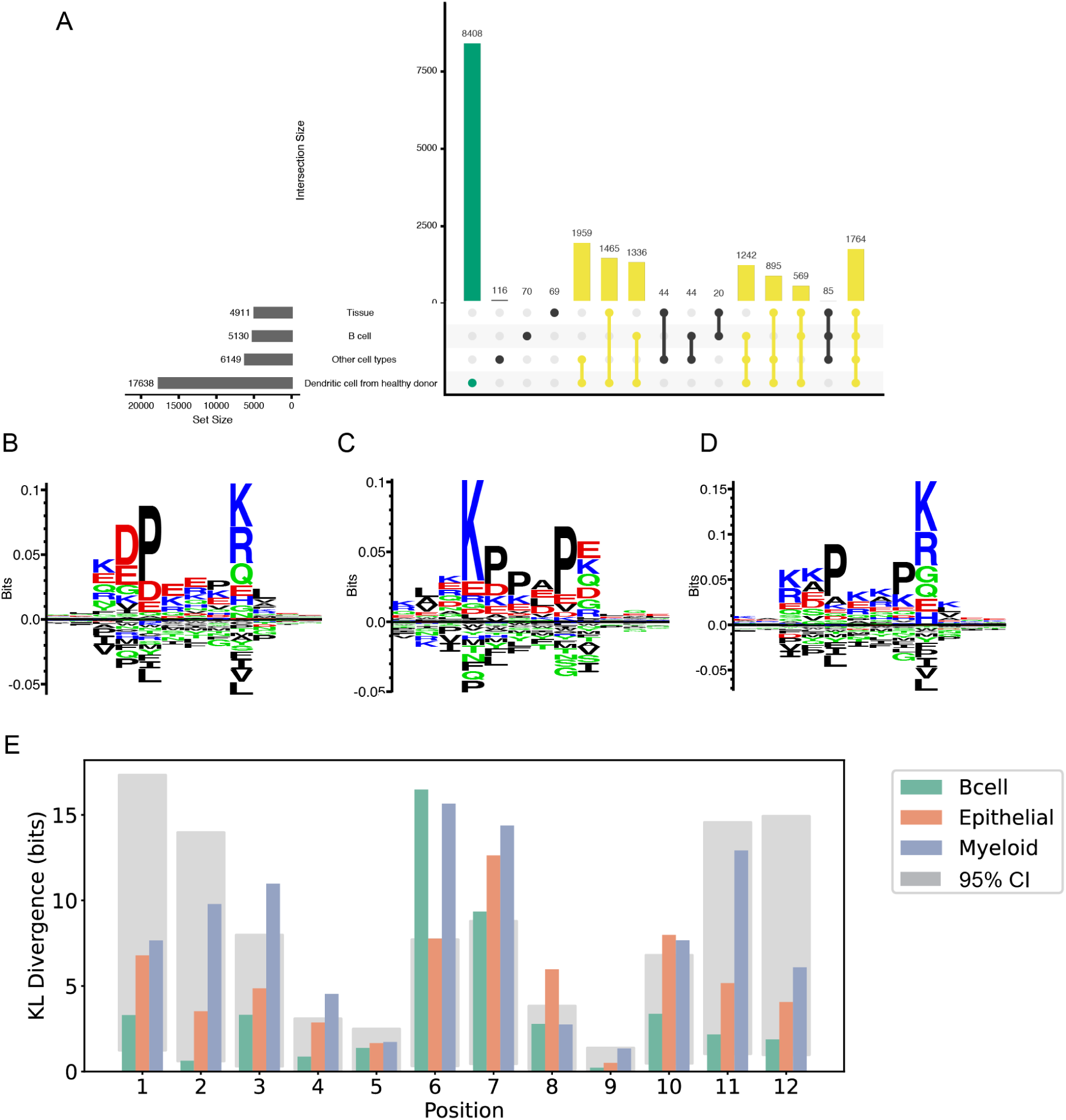
Comparison of antigen-processing signals across antigen-presenting cell types. A) Comparison of the proteins presented by dendritic cells (DCs) from healthy donors with other MHC II immunopeptidomics datasets available in IEDB. Green bars indicate proteins unique to the DC dataset, yellow bars indicate proteins shared between DCs and other cell types, and black bars indicate proteins detected only in other studies. The large overlap demonstrates that the DC dataset captures the majority of the known healthy human immunopeptidome. B-D) Sequence logos show the p-weighted Kullback–Leibler (KL) context motifs for peptides eluted from B cells, myeloid cells, and epithelial cells. E) The barplot shows the measured KL divergence between the probability-weighted Kullback-Leibler matrices of the three APC types. The shaded areas indicate 95% confidence intervals. The motifs are largely conserved across cell types, with no statistically significant differences (adjusted p > 0.05) in amino acid distributions compared with dendritic cells.

Overall, only a limited number of proteins (n = ≈450, representing the summed non-overlapping categories in the plot) were uniquely detected in other cell types but absent from our DC dataset. Nearly all proteins previously observed in healthy human immunopeptidomics studies were represented among the peptides presented by our DCs. We therefore conclude that our DC dataset provides comprehensive coverage of the proteins naturally presented on HLA-DR in healthy individuals and is suitable for comparative analyses with the autoimmune datasets.

We next tested whether differences in antigen-processing enzymes between APC types could account for the divergent sequence motifs observed in healthy donors, RA, and MS patients. Peptides and their immediate flanking regions (three residues upstream and downstream) were retrieved from IEDB for three major APC groups, respectively B cells, myeloid cells, and epithelial cells, yielding 47,964, 46,597, and 11,611 peptides. We generated p-weighted Kullback–Leibler (KL) sequence logos for each group and compared their flanking sequence motifs to those of our healthy donor DC dataset (Fig. 6B-D, Fig.6E). The overall patterns were highly similar across cell types, including a consistent proline enrichment at position 5 and lysine at position 9, characteristic of endosomal protease preferences. Quantitative comparison using KL divergence confirmed that none of the amino acid distributions differed significantly from the DC reference after multiple testing correction.

Notably, the amino acid enrichment at position 9, which was highly distinctive in the MS dataset, was not significantly different from any other cell type in the IEDB data and fell within the expected confidence interval.

These findings indicate that the variation observed in autoimmune samples cannot be attributed solely to differences in the antigen-presenting cell type or its proteolytic machinery. The disease-associated shifts in antigen-processing motifs are therefore more likely to reflect altered processing pathways or inflammatory modulation rather than baseline cell-type variability.

Together, these analyses demonstrate that our healthy donor DC dataset provides broad immunopeptidome coverage and that antigen-processing signatures are consistent across major APC types. Consequently, the processing differences observed in MS and, to a lesser extent, RA are unlikely to result from cell-type bias and instead may reflect disease-related alterations in the MHC class II pathway.

### Relative solvent accessibility and secondary structure of presented peptides

To assess whether structural properties of peptides differ between autoimmune and healthy antigen presentation, we analyzed the relative solvent accessibility (RSA) of peptides eluted from MHC class II molecules in the MS, RA and healthy donor datasets. RSA values for each residue were predicted using AlphaFold (Jumper et al. 2021) structural models of the source proteins as we did not have the complete structure for all proteins. The median RSA across the residues comprising each peptide was used as a summary measure.

Comparison of peptide RSA distributions between disease and healthy datasets using a Wilcoxon rank-sum test revealed significantly higher RSA values in both MS and RA (p < 0.001 for each), Fig. 7. This indicates that peptides presented during autoimmunity tend to originate from more solvent-exposed, surface-accessible regions of proteins. Such a pattern suggests that antigen processing in autoimmune settings may be altered, favoring peptides derived from outer, flexible protein regions. This could reflect an inflammatory or stress-associated shift in antigen-presenting cells, such as enhanced cathepsin activity or accelerated protein degradation, which increases access to solvent-exposed segments. The RSA increase was consistent across peptides from both tolerant and cryptic protein classes, indicating a general shift in processing rather than a cryptic protein–specific effect.

**Fig. 7.**
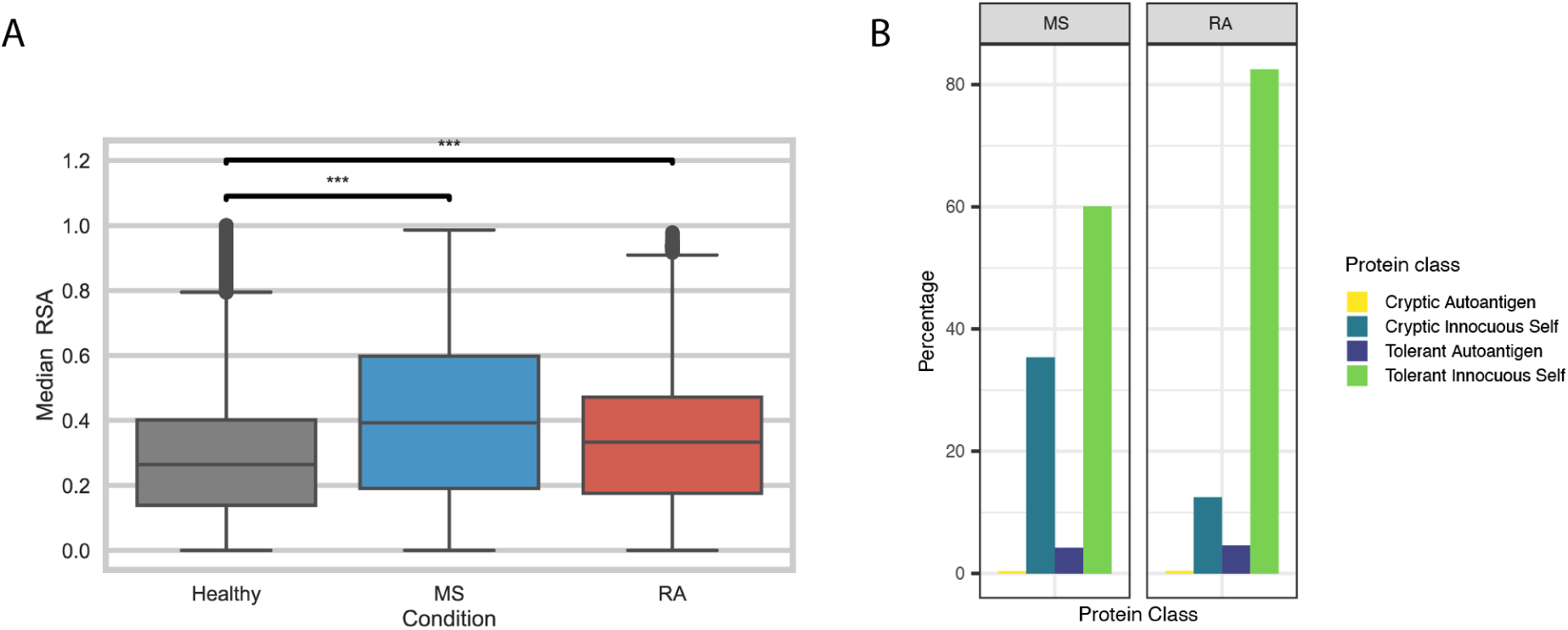
A) Median relative solvent accessibility (RSA) of MHC class II-bound peptides from healthy donors, rheumatoid arthritis (RA), and multiple sclerosis (MS) datasets. Peptides presented in autoimmune samples show significantly higher RSA values than healthy controls (p < 0.001, Wilcoxon test). B) The distribution (percentage per disease) of the proteins presented in the immunopeptidomics data of rheumatoid arthritis and multiple sclerosis patients classified into four types based on presentation in healthy donors and autoimmunogenic response by T cells.

We further compared the secondary structure composition of presented peptides among the datasets. Although several secondary-structure categories differed significantly between disease and healthy donors, all effect sizes were small, indicating that these differences are not biologically meaningful. Together, these findings point to a general shift in antigen processing in autoimmune disease toward presentation of peptides from more surface-exposed regions, rather than a structural bias specific to individual autoimmune proteins.

### Differential presentation of tolerant and cryptic proteins in autoimmune disease immunopeptidomes

To assess whether autoimmune diseases differ in their presentation of self-peptides, we analyzed the source proteins of MHC class II–bound peptides identified in immunopeptidomics datasets from patients with MS and RA. Source proteins were categorized as tolerant proteins also presented in healthy donors, or cryptic proteins uniquely presented in disease samples. The relative proportions of these classes are shown in Fig. 7B.

We expected that, despite disease status, the majority of presented peptides would originate from tolerant self-proteins. Indeed, this was the case for both diseases: most peptides were derived from tolerant proteins, consistent with normal presentation of self-antigens. However, we also observed a substantial fraction of peptides derived from cryptic proteins, particularly in MS. Among the 743 total proteins presented in MS, 268 (36%) were cryptic, compared with only 28 (12%) of 232 in RA. This enrichment of cryptic proteins in MS supports the hypothesis that altered antigen processing or presentation contributes to autoimmunity in organ-specific diseases, whereas systemic diseases such as RA may arise through other mechanisms, such as breakdown of tolerance or epitope spreading. Supplementary Material 1 S.Table 2 provides an overview of the proteins identified in the MS and RA datasets, categorized by disease association and protein classification.

To explore whether the increased presence of cryptic proteins could be linked to their cellular origin, we analyzed their predicted subcellular localization (Supplementary Material 4). On average, the distribution of cryptic proteins in MS and RA followed that expected by random sampling from the proteome. However, consistent with other cryptic innocuous self-proteins, they were slightly overrepresented among nuclear proteins. This supports that these proteins are normally inaccessible to MHC class II presentation in healthy individuals, and their appearance in disease likely reflects altered antigen processing, increased cell turnover, or release of intracellular material under inflammatory stress. The specific reasons why these particular cryptic proteins, and not others, become presented remain unclear. Future studies may focus on those cryptic proteins identified in extracellular or membrane compartments, as these could represent novel and accessible T cell autoantigens in MS and RA.

We next examined the presence of known autoantigens within each dataset. As expected, both diseases contained a mixture of tolerant and cryptic autoantigens (Fig. 4). Each disease also showed one cryptic autoantigen corresponding to its own pathology, fibrinogen beta chain (FGB) in RA (So et al. 2003) and myelin basic protein (MBP) in MS (Martinsen and Kursula 2022). Interestingly, the MS dataset also included peptides from the cryptic autoantigens FGB and SNRPD1, previously associated with RA and systemic lupus erythematosus (SLE), respectively, indicating potential overlap in autoantigen presentation across autoimmune phenotypes.

Both diseases also presented peptides derived from tolerant autoantigens, several of which overlapped between RA and MS. These included proteins such as ENO1, GSN, and HNRNPA2B1, which are known targets in multiple autoimmune conditions. Many of the shared tolerant autoantigens are associated with systemic diseases, such as Behçet’s disease, lupus, rheumatoid arthritis, and vasculitis, indicating that tolerant autoantigens may underlie shared immune activation across autoimmune phenotypes.

In summary, MS and RA immunopeptidomes display distinct proportions and types of self-protein presentation. MS exhibits a marked enrichment of cryptic proteins, largely nuclear and not normally accessible to MHC II processing, suggesting a broader disturbance in antigen processing or cellular integrity. In contrast, RA maintains a predominance of tolerant autoantigens, reflecting immune reactivity toward normally presented self-proteins and supporting different underlying mechanisms of autoimmunity between organ-specific and systemic diseases.

### Peptide placement in tolerant autoantigens

To determine whether disease-associated presentation of tolerant autoantigens involves novel epitopes or just reflects normal antigen display, we compared the peptides detected in RA and MS patients with those presented in healthy donors. In this analysis we specifically investigate the proteins containing autoreactive T cell epitopes in RA or MS based on the experimental data from IEDB.

Peptides from healthy donors, RA, and MS patients and known autoimmunogenic epitopes (tested in RA and MS patients; IEDB) were mapped to their corresponding source protein sequences. Representative peptide distributions for selected tolerant autoantigens are shown in Fig. 8, with the complete set presented in Supplementary Material 1 S.Fig. 3.

**Fig. 8.**
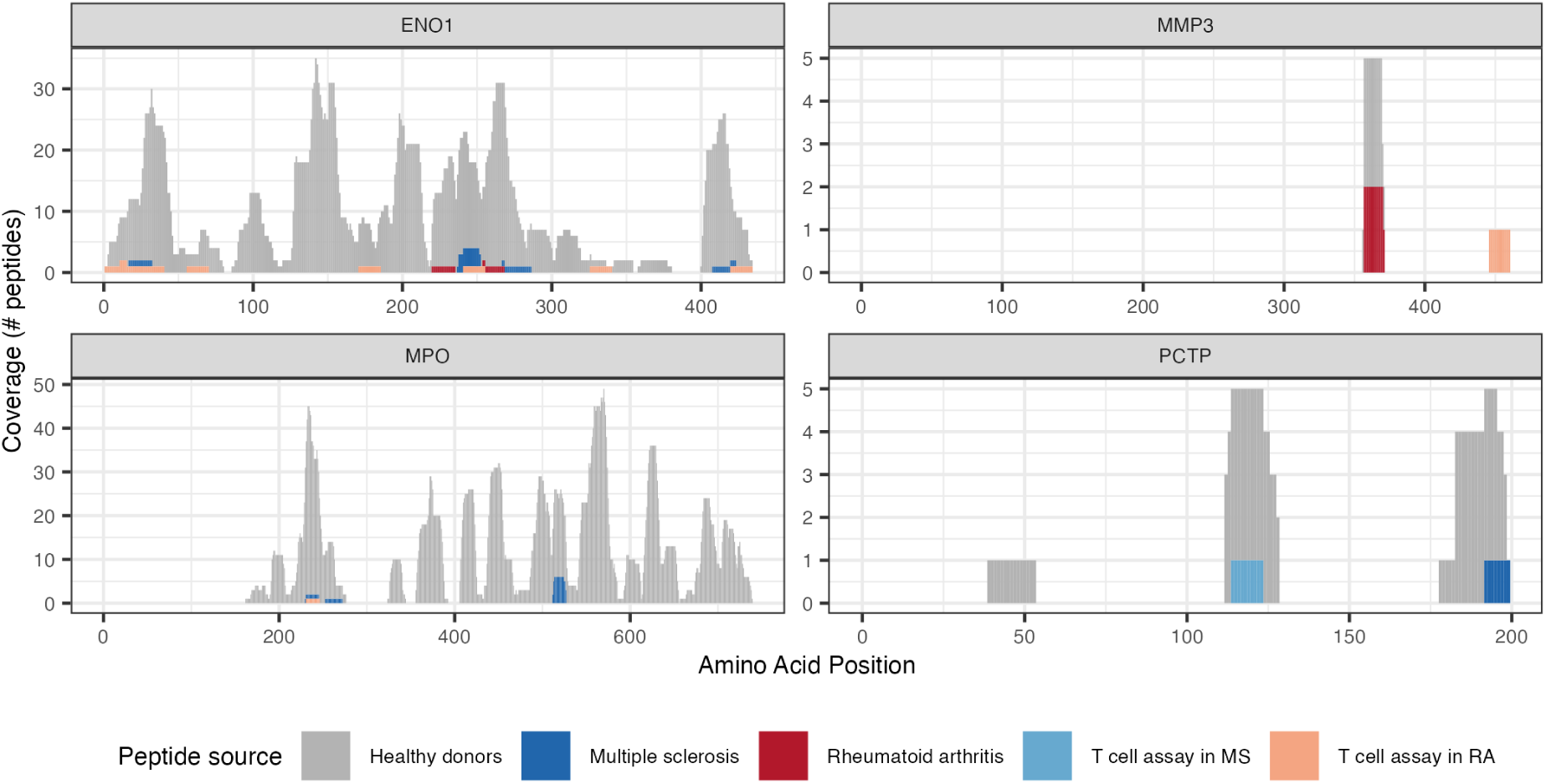
A selection of four tolerant autoantigens of multiple sclerosis and rheumatoid arthritis. Each plot is a protein with the sequence length on the x-axis. The peptides presented in the healthy donors are mapped in grey, the peptides presented by autoimmune patients are blue (MS) and red (RA). The coral and light-blue indicate autoimmunogenic epitopes from T cell assays based on RA patients and MS patients respectively.

Overall, the epitopes of tolerant autoantigens did not differ substantially from those presented in healthy donors, indicating that disease-associated presentation in RA and MS is not primarily driven by the emergence of novel epitopes. Instead, these findings suggest that autoimmunity involving tolerant autoantigens arises through mechanisms such as molecular mimicry, epitope spreading, or breakdown of peripheral tolerance.

In RA, we observed a high degree of concordance between peptides presented in patients and those found in healthy donors. For example, ENO1 peptides spanned the full protein sequence in both groups, and similar overlaps were observed for ANXA2, CALR, FLNA, GSN, HNRNPA2B1, TNC, and VIM. For MMP3, only a restricted region of the protein was detected in healthy donors, yet this same segment was also presented in RA. A few exceptions, such as FGA and GNS, exhibited limited overlap, with RA-derived peptides mapping to slightly different regions.

A similar pattern was observed in MS, where the majority of tolerant autoantigens, such as ACTB, CD74, ENO1, GSN, H2BC4, H4C1, HLA-DRA, HSPA5, HSPB1, MPO, and RPL7, showed extensive or complete overlap with peptides from healthy donors. Minor deviations were noted for proteins like PCTP (near-complete overlap), H2BC11 (presentation of distinct distal regions in MS), and HNRNPA2B1, where MS-derived peptides included one site not detected in healthy donors (residues 291–306) and another partially overlapping with immunogenic epitopes from IEDB. FGA and HLA-DRB5 also displayed only partial overlap.

Given the extensive overlap observed, we did not pursue further analyses of peptide placement within secondary or tertiary structures, as the data did not indicate a structural contribution to altered antigen presentation.

## Discussion

Here, we combined large-scale characterization of known autoantigens with healthy donors immunopeptidomics, and demonstrated that autoantigens segregated into two mechanistic classes, tolerant and cryptic, depending on whether their source proteins were naturally presented under physiological conditions. This empirical distinction uncovered consistent molecular and functional differences between the two groups. Tolerant autoantigens were predominantly extracellular and enriched for immune-related functions, whereas cryptic autoantigens were overrepresented among membrane-associated and lysosomal/vacuolar proteins, and closely linked to disease-specific cellular processes. These observations suggest that not all autoantigens arise through the same pathogenic route. We therefore propose two complementary mechanisms leading to autoimmune recognition: one driven by *loss of tolerance* toward normally presented self-proteins, and another arising from *altered antigen processing and presentation* that exposes previously unpresented, cryptic self-peptides. Central tolerance, governed by AIRE-dependent antigen presentation in the thymus, eliminates T cells reactive to peptides that are normally displayed during development. Self tolerance therefore, can be interpreted as tolerance to the “presented” self. Cryptic autoantigens, or self peptides that are not naturally presented, would then be likely to escape central tolerance, and remain unseen by reactive T cells until alternate processing exposes them later in life. In contrast, tolerant autoantigens would be regularly presented but would become reactive when peripheral tolerance mechanisms fail. For instance failure of regulatory T cell suppression during inflammation or immune dysregulation.

Evidence supporting this dual-path model comes from our immunopeptidomic analyses of RA and MS patients, which revealed distinct antigen-processing signatures compared to healthy donors. Both autoimmune datasets contained peptides from cryptic proteins which could not be explained by differences of the cell types the experiments were conducted in. In particular, MS displayed altered proteolytic cut-site preferences, suggesting an antigen-processing change that may facilitate presentation of cryptic peptides. Furthermore, the peptides from the autoimmune patients had significantly higher relative solvent accessibility values also pointing towards a change in the antigen processing milieu.

We propose that in the case of tolerant autoantigens, autoimmune activation results from peripheral tolerance breakdown through mechanisms such as molecular mimicry or epitope spreading. Several autoimmune diseases have been associated with pathogen-driven molecular mimicry. Epstein–Barr virus infection for instance has been linked to the onset of MS (Bjornevik et al. 2022), where immune responses initially directed toward a pathogen cross-react with structurally similar self-epitopes. Epitope spreading, in turn, can propagate these responses as new self-epitopes become exposed during tissue inflammation (Boldison et al. 2015; Vanderlugt and Miller 2002). The resulting immune activation could further diversify the targets of the immune response, enhancing presentation of tolerant autoantigens at extracellular sites and perpetuating autoimmunity. Conversely, cryptic autoantigens may emerge when inflammation or cellular stress disrupts normal antigen processing, leading to the display of self-peptides that are usually hidden from immune surveillance. It has previously been proposed, in the context of post-translational modifications, that alterations of the presented peptidome can transform tolerance to self to autoreactivity (Lichti and Wan 2023). The underlying causes of such antigen-processing alterations remain uncertain but may involve changes in protease activity, peptide editing, or altered endosomal trafficking.

Together, our results suggest that *loss of tolerance* and *antigen processing and presentation alterations* represent complementary, rather than competing, mechanisms in the development of autoimmunity, offering a mechanistic framework that reconciles diverse autoimmune phenotypes.

We observed that most autoimmune diseases were dominated by a single class of autoantigens, with cryptic autoantigens significantly enriched in organ-specific diseases and tolerant autoantigens prevailing in systemic diseases. Although the classification of autoimmune diseases into organ-specific and systemic categories is widely used, it remains largely descriptive rather than mechanistic. Both groups involve overlapping immune processes, including autoreactive T and B cells, cytokine networks, and complement activation, and there is no consensus on what defines an organ-specific or systemic disorder. Rheumatoid arthritis (RA) exhibits systemic autoantibody responses (ACPA, RF) and extra-articular symptoms, but primarily affects the joints leading some to classify it as organ-specific rather than systemic (Ostrov 2015; L. Wang et al. 2015, 201). Thus, while this dichotomy provides a useful clinical framework, it may oversimplify the complex immunopathology underlying autoantigen presentation and disease manifestation.

Our immunopeptidomics analyses indicate that antigen processing is altered in autoimmune disease, particularly in multiple sclerosis (MS) and, to a lesser extent, rheumatoid arthritis (RA). Both diseases exhibited distinct proteolytic signatures, differences in the surface accessibility of presented peptides, and evidence of presentation of proteins that are normally absent from the healthy MHC class II immunopeptidome.

The detection of such cryptic proteins suggests that antigen-presenting cells (APCs) in autoimmune settings process a broader or atypical pool of self-proteins. This could arise from increased proteolytic activity or altered intracellular trafficking during inflammation. Cytokine exposure, oxidative stress, or metabolic activation of APCs can modulate endosomal acidification and protease expression, particularly of cathepsins, thereby changing which proteins are degraded and loaded onto MHC class II molecules (Honey et al. 2001; Crotzer et al. 2010; Laha et al. 1995). Such changes could render previously inaccessible, cryptic epitopes available for presentation and T cell recognition. At the peptide level, we observed clear alterations in terminal amino acid preferences in disease, with reduced proline dependency and an increased presence of hydrophobic or basic residues in MS peptides. These findings are consistent with a shift in cathepsin activity or altered regulation of peptide editors such as HLA-DM and HLA-DO, which could modify the trimming or selection of bound peptides (Denzin 2013). Finally, peptides eluted from autoimmune samples displayed higher relative surface accessibility (RSA) compared to those from healthy donors, suggesting that disease-associated antigen processing may preferentially generate peptides from more exposed protein regions. Together, these results support a model where inflammatory or stress-induced changes in APC processing pathways expand the repertoire of presented self-peptides, thereby increasing the likelihood of cryptic epitope presentation and autoimmune activation. Several antigen processing steps have previously been associated with autoimmune diseases. For instance, altered cathepsin activity that might be caused by change in endosomal pH, redox balance, or cytokine-mediated signaling, could lead to non-canonical cleavage events that generate cryptic peptides. Specifically, Cathepsin V has been seen to be overexpressed in myasthenia gravis (Tolosa et al. 2003), and other studies have implicated cathepsins S and B in shaping the CD4⁺ T cell repertoire through selective cleavage patterns (Høglund et al. 2020).

HLA-DM and HLA-DO regulate peptide editing and exchange by removing low-affinity peptides and mediating CLIP release from MHC class II molecules. Subtle changes in the DO:DM ratio can substantially change the immunopeptidome. A higher amount of free DM will result in a more narrow peptide pool with stronger MHC binding affinity (Olsson et al. 2022; Nanaware et al. 2019). Experimental modulation of DO expression has demonstrated that reduced DO enhances the presentation of low-affinity peptides (Welsh et al. 2020). Dysregulation of this system has been observed in an autoimmune context. For instance, in type 1 diabetes, higher MHCII-CLIP levels were detected despite normal DM/DO expression ratios, suggesting altered functional dynamics of peptide loading (Gilles et al. 2023). Such changes could reduce editing stringency, allowing unstable or low-affinity peptides to be presented on the cell surface.

Together, our findings support a model in which autoimmunity arises not only from the identity of self-proteins but also from how the antigen-processing machinery handles them. Altered proteolytic activity, peptide editing, or endosomal conditions could expand the pool of peptides loaded onto MHC class II molecules, thereby exposing cryptic epitopes that normally escape immune surveillance. This may explain the higher proportion of cryptic peptide presentation observed in multiple sclerosis compared with rheumatoid arthritis, where antigen-processing appears more conserved. Defining autoantigens as either tolerant or cryptic highlights two complementary pathways toward loss of self-tolerance: one rooted in regulatory failure of the immune system, and another in altered antigen processing. These mechanistic classes show distinct molecular features, reflecting their differential access to the MHC class II pathway. Recognizing this dichotomy provides a biological framework that connects antigen presentation to autoimmune diversity and suggests that altered processing, rather than antigen availability alone, may be a decisive factor in triggering pathogenic T cell responses.

Recognizing these distinct pathways has direct implications for biomarker discovery and therapeutic design, as altered antigen processing may serve as a measurable feature of immune dysregulation. Integrating immunopeptidomics with molecular profiling in patient samples could therefore pave the way for precision diagnostics and interventions that target the early stages of autoimmune activation.

## Contributors

CBQ and ABS conceived and designed the study. ABS collected, analyzed and interpreted the data. SRA created the original analysis on cell type processing, later modified by ABS. FDH created the nested MHC enrichment scores, and extracted data from alphafold. ABS wrote the first draft of the manuscript, and CBQ edited and reviewed the manuscript. All authors read and approved the final manuscript.

## Acknowledgements

This research has been supported by the Technical University of Denmark (DTU), Department of Health Technology.

Graphical abstract created in BioRender. Saksager, A. B. (2026) https://BioRender.com/vjgfy80

## Data Sharing Statement

Data sharing is not applicable to this article as no datasets were generated during the current study.

## Declaration of generative AI and AI-assisted technologies in the manuscript preparation process

During the preparation of this work the authors used ChatGPT in order to paraphrase, rephrase, or improve the clarity and readability of text originally written by the author. After using this tool, the authors reviewed and edited the content as needed and take full responsibility for the content of the published article.

