## Supplementary Material 1 for "Antigen-processing rewiring expose cryptic self promoting organ-specific autoimmunity"

### Supplementary Information

#### Content

Classification of autoimmune diseases: Supplementary Table 1……………………………………………….1
Presented Proteins on MHC class II in Rheumatoid arthritis and Multiple Sclerosis patients: Supplementary Table 2 …………………………………………………………………………………………………….2
Significantly different physico-chemical protein features: Supplementary Fig.1…………………………….....3
Position-specific Sequence Matrices: Supplementary Fig. 2…………………………………………………...4
Tolerant autoantigens in rheumatoid arthritis and multiple sclerosis: Supplementary Fig. 3……………………..5
References ..…………………………………………………………………………………………………6

### Classification of autoimmune diseases

**Supplementary Table 1: Organ-specific and systemic classifications of autoimmune diseases.**

| Disease | Type | Organs | Papers | Paper with other classification |
|---|---|---|---|---|
| ORGAN-SPECIFIC | | | | |
| Alopecia areta | Organ-specific | Hair follicle | (Trüeb and Dias 2018; Deng et al. 2024) | |
| Autoimmune hepatitis | Organ-specific | Liver | (Wang et al. 2015; Longhi et al. 2025) | |
| Autoimmune thyroditis | Organ-specific | Thyroid | (Mastrandrea 2015; Vargas-Uricoechea 2023) | |
| autoimmune uveitis | Organ-specific | Eye | (Papotto et al. 2014; Thurau and Wildner 2003) | |
| Goodpasture syndrome | Organ-specific | Kidney/Lung | (Mastrandrea 2015) | |
| Graves | Organ-specific | Thyroid | (Mastrandrea 2015, Wang et al. 2015) | |
| Multiple sclerosis | Organ-specific | Central nervous system | (Harris 2016, Wang et al. 2015) | |
| Myathenia gravis | Organ-specific | Muscle | (Mastrandrea 2015) | |
| Pemphigus | Organ-specific | Skin | (Kridin 2018) | |
| Primary biliray cholangitis | Organ-specific | Bile ducts | (Wang et al. 2015) | |
| Type 1 diabtetes melitus | Organ-specific | Pancreatic islets | (Mastrandrea 2015, Wang et al. 2015) | |
| Vitiligo | Organ-specific | Skin | (Harris 2016) | |
| SYSTEMIC DISEASES | | | | |
| Autoimmune vasculitis | Systemic | kidneys, eyes, sinuses, peripheral nerves, skin, and upper and lower respiratory tracts. | (Sharma et al. 2024) | |
| Psoriasis | Systemic | systemic, skin | (Hao et al. 2021; Bu et al. 2022) | |
| Rheumatoid arthritis | Systemic | Systemic: joints, surrounding bone, blood vessels, and connective tissue, lungs, heart, eyes, skin, and various other body systems. | (Ostrov 2015) | (Wang et al. 2015) |
| Sjogren's syndrome | Systemic | Several organs (e.g. lungs, liver, kidneys, central nervous system), mainly salivary and lacrimal glands | (Ostrov 2015, Wang et al. 2015) | |
| Systemic lupus erythematosus | Systemic | Several organs (heart, joints, skin, lungs, blood vessels, liver, kidneys, and nervous system) | (Ostrov 2015, Wang et al. 2015) | |
| Vog-koyanagi-Harada disease | Systemic | Multiple systems incl. nervous system and skin | (Greco et al. 2013; Sakata et al. 2014) | |
| Bechets disease | Systemic | Systemic, skin, eyes, joints | Mendoza-pinto2010 | |

| Disease | Type | Organs | Papers | Paper with other classification |
|---|---|---|---|---|
| | | | (Mendoza-Pinto et al. 2010) | |
| UNCLASSIFIED | | | | |
| Nueromyelitis optica | Conflicting, lacking | | | Samim2024 |

**Supplementary Table 2: Classification of presented proteins on MHC class II in rheumatoid arthritis patients and multiple sclerosis patients, according to the presentation in healthy donors.**

| Disease | Total number of unique proteins presented | Cryptic IEDB disease-specific autoantigens | Cryptic IEDB other disease autoantigens | Cryptic proteins | Tolerant IEDB disease-specific autoantigens | Tolerant IEDB other disease autoantigen |
|---|---|---|---|---|---|---|
| Rheumatoid Arthritis | 232 | FGB | - | 28 proteins | FGA, GSN, ENO1, MMP3, HNRNPA2B1, TNC, MYH9, CHI3L1 | HSPD1 (Bechets, MS, T1D), HLA-DRA, HLA-DRB5 (MS) , CD74 (gravis) |
| Multiple Sclerosis | 743 | MBP | FGB (RA), SNRPD1 (RA, Lupus) | 268 proteins | HLA-DRA, HLA-DRB5, PCTP | FGA (RA)<br>CD74 (gravis)<br>H2BC4 (RA)<br>ENO1 (RA)<br>H2* (lupus)<br>MPO (vasculitis, RA)<br>H4* (RA, lupus)<br>GSN (RA)<br>HNRNPA2B1 (RA)<br>RPL7 (lupus) |

Significantly different physico-chemical protein features

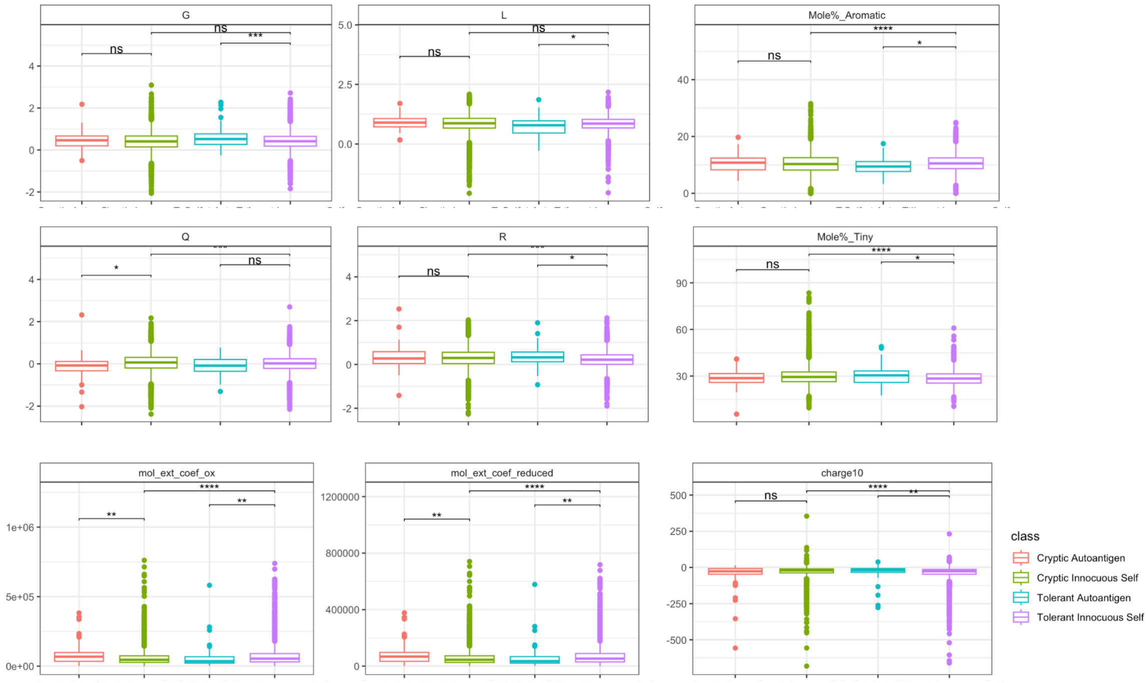


**Supplementary Fig. 1.** Selected amino acid compositions and physico-chemical features in which either the two tolerant or the two cryptic groups of proteins are significantly different from each other. The significance level between the two groups of innocuous self is shown, as well as the comparison between the two tolerant groups and two cryptic groups. The asterisks indicate * = 0.05, ** = 0.01, *** = 0.001

Position-specific Sequence Matrices

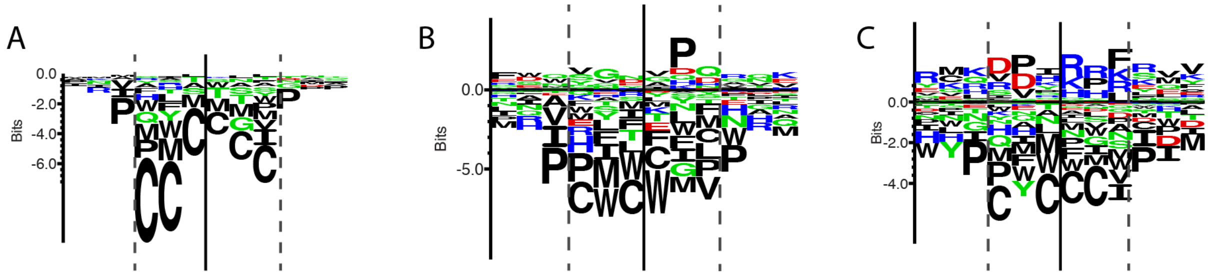


**Supplementary Fig. 2.** Position-specific sequence matrices of the context of A) healthy donors peptides B) rheumatoid arthritis patients peptides, and C) multiple sclerosis patients peptides.

Tolerant autoantigens in rheumatoid arthritis and multiple sclerosis

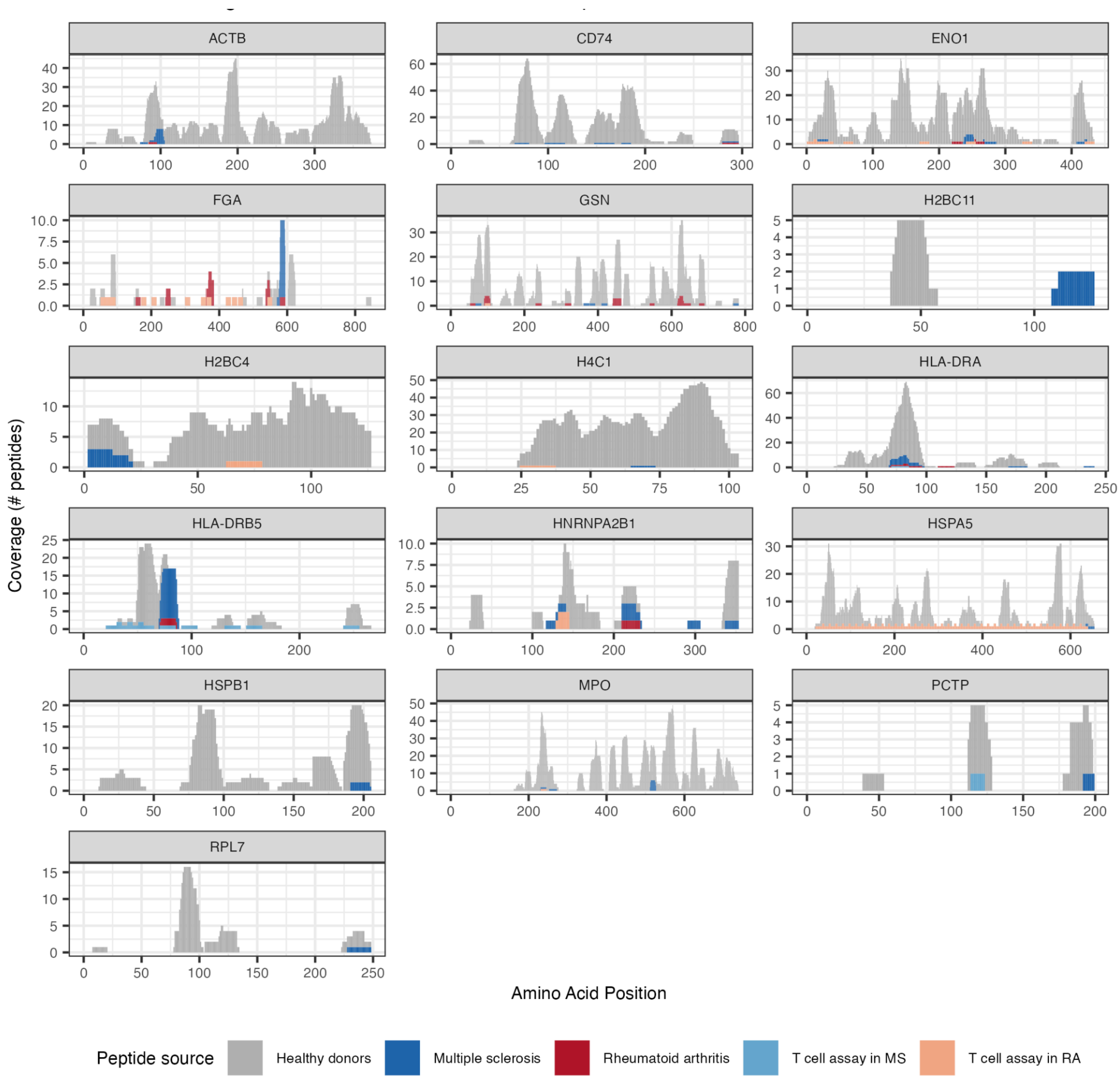


**Supplementary Fig. 3.** All tolerant autoantigens of multiple sclerosis and rheumatoid arthritis. Each plot is a protein with the sequence length on the x-axis. The peptides presented in the healthy donors are mapped in grey, the peptides presented by autoimmune patients are blue (MS) and red (RA). The coral and light-blue indicate autoimmunogenic epitopes from T cell assays based on RA patients and MS patients respectively. If they do not have any T cell assays indicated here, it means that the T cell assay classifying the autoantigen were tested in another autoimmune disease.
